# A high-quality phased genome assembly of stinging nettle, *Urtica dioica* ssp. *dioica*

**DOI:** 10.1101/2024.11.21.624787

**Authors:** Kaede Hirabayashi, Christopher R. Dumigan, Matúš Ku•ka, Diana M. Percy, Gea Guerriero, Quentin Cronk, Michael K. Deyholos, Marco Todesco

## Abstract

Stinging nettles (*Urtica dioica*) have a long history of association with human civilization, having been used as a source of textile fibres, food and medicine. Here, we present a chromosome-level, phased genome assembly for a diploid female clone of *Urtica dioica* from Romania. Using a combination of PacBio HiFi, Oxford Nanopore and Illumina sequencing, and Hi-C long-range interaction data (using a novel Hi-C protocol presented here), we assembled two haplotypes of 574.9 Mbp (contig N50 = 10.9 Mbp, scaffold N50 = 44.0 Mbp) and 521.2 Mbp (contig N50 = 13.5 Mbp, scaffold N50 = 48.0 Mbp), with assembly BUSCO scores of 92.6% and 92.3%. We annotated 20,333 and 20,140 genes for each haplotype, covering over 90% of the complete BUSCO genes and including two copies of a gene putatively encoding the neurotoxic peptide urthionin, which could contribute to nettle’s characteristic sting. Despite its relatively small size, the nettle genome displays very high levels of repetitiveness, with transposable elements comprising more than 60% of the genome, as well as considerable structural variation. This genome assembly represents an important resource for the nettle community and will enable the investigation of the genetic basis of the many interesting characteristics of this species.

## 1. Introduction

*Urtica dioica* L. (*U. dioica* ssp. *dioica*; stinging nettle, or common nettle) is an herbaceous perennial in the Urticaceae family, native to Eurasia and northwest Africa [1]. *U. dioica* is widely distributed in temperate and tropical climates and can grow on a variety of different soil types, although it prefers moist habitats rich in nitrogen and phosphorus [2]. It is often found on disturbed soils, where it can grow in dense stands, which has earned *U. dioica* a reputation as a weed. However, since at least the Bronze Age and continuing to the present day, *U. dioica* and its close relatives have also been used by humans as sources of fibre, food, and medicine [3–5]. Like cannabis (*Cannabis sativa* L.), flax (*Linum usitatissimum* L.), and ramie (*Boehmeria nivea* (L.) Gaudich.), the stems of *U. dioica* produce long (43-58 mm) bast fibres that are rich in crystalline cellulose (>70%) and have a relatively low lignin content (2-7%) [6]. Fibres have been used historically in papermaking, cordage, and textiles, and there is renewed interest in using nettle in both clothing and composite materials. Both the shoots and roots of stinging nettle are consumed in some cultures, and dried nettle powder contains an impressive 30% protein content (w/w) [7]. The lipids produced in the leaves also contain a very high proportion of polyunsaturated fatty acids (41% alpha-linolenic acid, 18:3n-3; 12% linoleic acid, 18:2n-6), which are beneficial to human nutrition [8]. Teas and other extracts made from stinging nettle have been reported to have anti-proliferative, antibacterial, anti-inflammatory, hypoglycemic, and other pharmacological activities [9].

The common name of stinging nettle comes from the irritation experienced by vertebrates when they make skin contact with certain trichomes of *U. dioica*. Although this effect was previously attributed to the presence of histamines, organic acids, and neurotransmitters [10], it has recently been demonstrated that small peptides, namely urthionin (•-Uf1a) and a sodium-gated channel modulator named urticatoxin (•/•-Uf2a), contribute to the painful sensation [11]. Small neurotoxic peptides in *Dendrocnide* species similarly affect sodium-gated channels [12].

*U. dioica* is either diploid (2n = 24, 26) or tetraploid (2n = 48, 52), with tetraploids being much more abundant [2,13]. Occasionally, triploid and pentaploid individuals are found within populations [2]. Genome size estimates for diploid *U. dioica* also vary according to different studies, ranging from 558 Mbp to 660 Mbp [2,13]. *U. dioica* is distinct from slender nettle, *Urtica gracilis* Aiton (also known as *U. dioica* ssp. *gracilis* (Aiton) Selander), which is native to North America. While *U. dioica* is generally dioecious, meaning that female and male flowers are on separate individuals, *U. gracilis* is monoecious with female and male flowers on the same plant [14]. Complex variation of morphological traits in the genus Urtica makes the classification of species and subspecies sometimes difficult, although attempts have been made to resolve these phylogenetic relationships using molecular data [15]. A high-quality reference genome of a well-characterized *U. dioica* individual would greatly facilitate the study of the phenotypic, genetic and karyotypic diversity within *U. dioica* and help improve understanding of taxonomic relationships within the Urticaceae.

In this study, we develop a diploid, phased chromosome-level genome assembly for *Urtica dioica* ssp. *dioica* using a combination of sequencing approaches, including PacBio HiFi and Oxford Nanopore (ONT) long reads, whole genome shotgun (WGS) Illumina short reads and chromatin conformation capture (Hi-C). We further explore characteristics of the stinging nettle genome, such as the repeat landscape across chromosomes, the presence of large structural variants (SVs) between haplotypes, and how it compares to related taxa. Finally, we comment on the presence of putative neurotoxic peptides associated with the nettle’s sting.

## 2. Results and Discussion

### 2.1 Genome sequencing and assembly

We generated 66.3 Gbp of PacBio HiFi data (115X genome coverage) from a female, diploid individual of stinging nettle. HiFi reads had a median read length of 15.45 kbp and a median read quality of Q29. Only reads with quality •Q20 were used for the assembly (50.7 Gbp, 88X genomic coverage). We also generated 29.02 Gbp paired-end Hi-C reads (50X genome coverage), with 96.49% of the reads with quality >Q30. The combination of long reads and long-range interaction data produced an initial phased contig-level assembly. Haplotype 1 (H1) had a total length of 574.795 Mbp and contig N50 = 20,59 Mbp (1,404 contigs); haplotype 2 (H2) had a total length of 521,297 Mbp and N50 = 24.787 Mbp (229 contigs). There is a considerable range of genome size estimates for *Urtica dioica* from previous studies [2,13], so we performed confirmatory flow cytometry on the individual that we sequenced, which was originally estimated at 1C = 650 Mb [13]. Our new estimate, reported here, is 2C = 1.26 pg, which translates to 1C = 616 Mbp. While our assembled genome is smaller than that estimate, it has high BUSCO completeness scores (>92% for each haplotype) and kmer completeness (98.15% for both haplotypes combined; Table S1, Figure S1), consistent with recent suggestions that flow cytometry over-estimates genome sizes [16]. Scaffolding using Hi-C data anchored contigs into 13 pseudochromosomes for each haplotype. We visually inspected Hi-C contact maps for the two haplotypes and manually corrected misjoins and misassemblies, retaining only putative structural variants (SVs) whose presence was strongly supported by Hi-C data and by the presence of long HiFi and ONT reads spanning their borders. Finally, we compared the chromosome organization of the haplotype assemblies to that of the *U. dioica* assembly produced by the Darwin Tree of Life project [17] to ensure that Hi-C contact maps supported our contig ordering (Figure S2). The final H1 assembly had a total length of 574.93 Mbp, scaffold N50 = 43.96 Mbp, and contig N50 = 10.89 Mbp, where 92.59% of the contigs were above 50 kbp. The final H2 assembly was of similar quality, with a total length of 521.16 Mbp, scaffold N50 = 47.99 Mbp, and contig N50 = 13.53 Mbp with 98.98% of contigs above 50 kbp in length. BUSCO score was 92.6% complete for H1 and 92.3% complete for H2, with very low levels of duplication (<3%) (Table 1; Figure 1). While 89.5% and 97.8% of the H1 and H2 genome were placed in the chromosomes respectively, we had 72.33 Mbp of unplaced sequences. We note that 95.44% of these sequences (69.0 Mbp) were repetitive, and we only detected five BUSCO genes within them. Hifiasm placed all the repetitive sequences that could not be unequivocally assigned to either haplotype in H1, which explains the difference in size between H1 and H2.

**Table 1.**
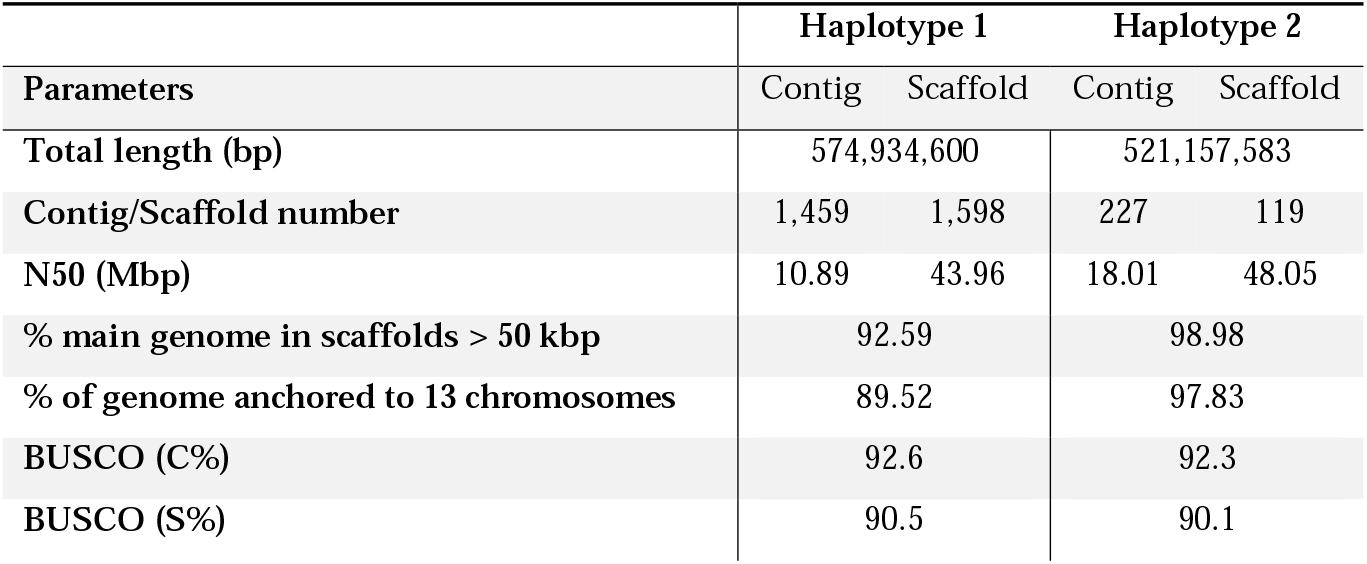
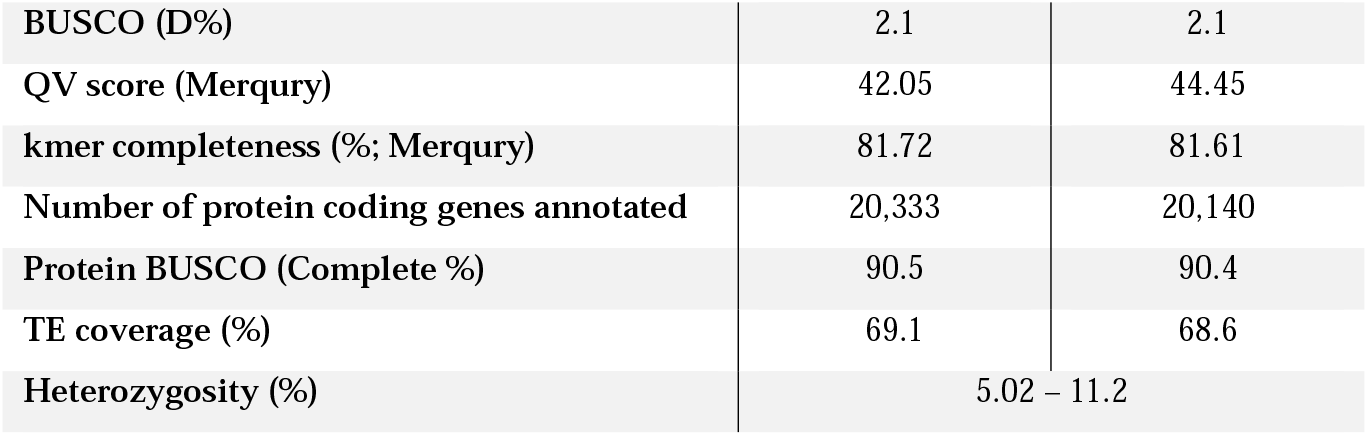
Genome assembly statistics.

| Parameters | Haplotype 1 |  | Haplotype 2 |  |
| --- | --- | --- | --- | --- |
|  | Contig | Scaffold | Contig | Scaffold |
| **Total length (bp)** | 574,934,600 |  | 521,157,583 |  |
| **Contig/Scaffold number** | 1,459 | 1,598 | 227 | 119 |
| **N50 (Mbp)** | 10.89 | 43.96 | 18.01 | 48.05 |
| **% main genome in scaffolds > 50 kbp** | 92.59 |  | 98.98 |  |
| **% of genome anchored to 13 chromosomes** | 89.52 |  | 97.83 |  |
| **BUSCO (C%)** | 92.6 |  | 92.3 |  |
| **BUSCO (S%)** | 90.5 |  | 90.1 |  |

|  |  |  |
| --- | --- | --- |
| BUSCO (D%) | 2.1 | 2.1 |
| QV score (Merqury) | 42.05 | 44.45 |
| kmer completeness (%; Merqury) | 81.72 | 81.61 |
| Number of protein coding genes annotated | 20,333 | 20,140 |
| Protein BUSCO (Complete %) | 90.5 | 90.4 |
| TE coverage (%) | 69.1 | 68.6 |
| Heterozygosity (%) | 5.02 – 11.2 |  |

**Figure 1.**
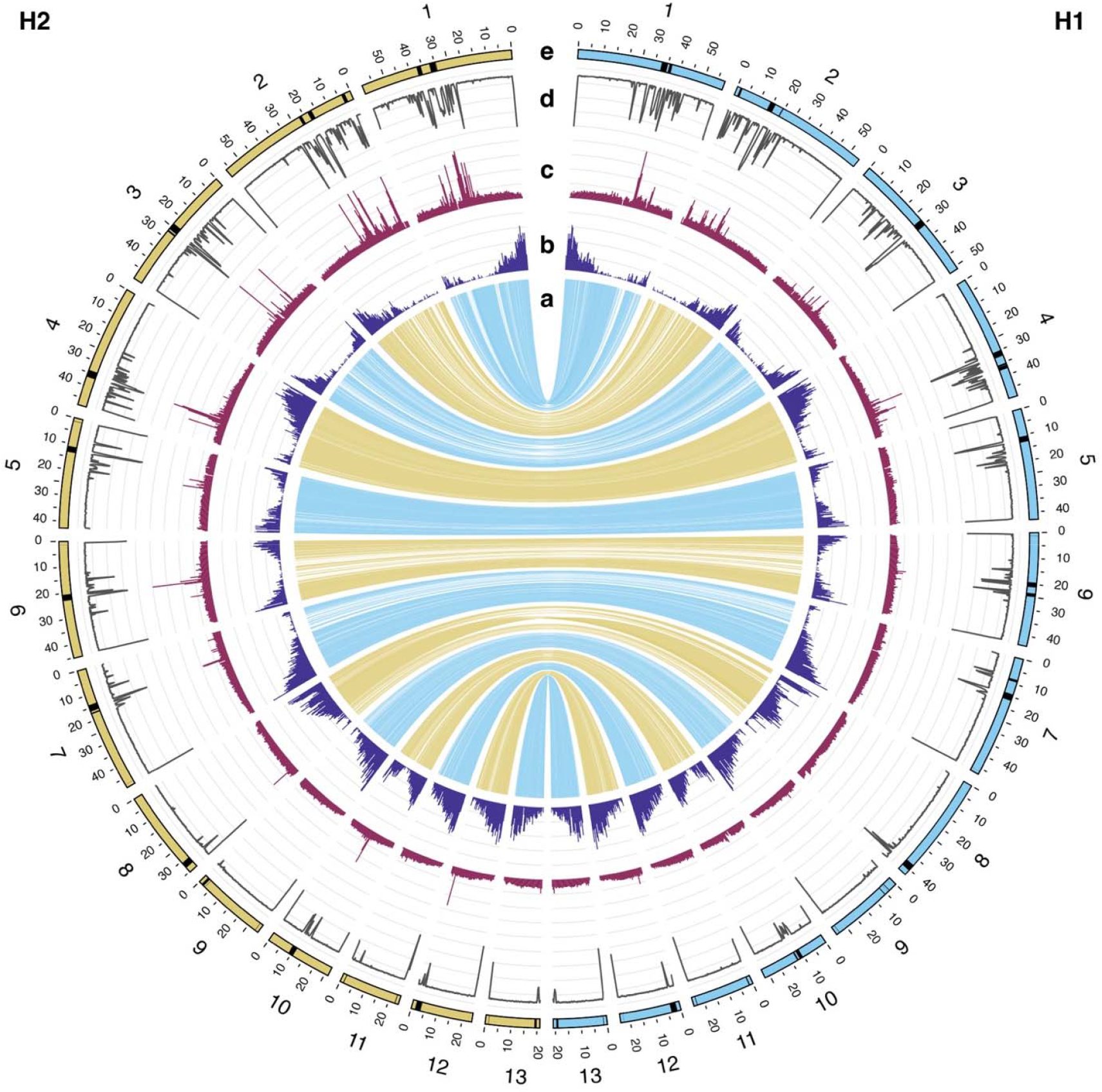
Haplotype-resolved assembly of a female *Urtica dioica* ssp. *dioica* individual (2n = 26). The two haplotypes are compared (H1 on the right, blue; H2 on the left, yellow). The tracks in the Circos plot represent (**a**) aligned regions between haplotypes, (**b**) gene density, (**c**) TE density, (**d**) repeats Shannon diversity score, and (**e**) predicted centromeric regions, highlighted in black on the ideogram.

### 2.2 Genome structure

We expected to find very high synteny between the two haplotypes of our stinging nettle assembly, especially since we had high HiFi long-reads and Hi-C coverage, and thorough manual curation should minimize the chances of misassemblies between haplotypes. However, we observed numerous structural variants (SVs) between haplotype assemblies (see chromosomes 1, 2, 3, and 8 in Figure 2, Figure S3). In particular, we found that the two haplotypes differed by a massive inversion (18.4 Mbp in length) on chromosome 8, which was surrounded by multiple duplicated regions (Figure 3). We manually checked all those SVs using Hi-C data and looked for HiFi and Nanopore reads spanning their breakpoints; this included manually “correcting” these SVs in either haplotype to determine if this resulted in improved Hi-C contact maps. In all cases, these checks supported the current chromosome organization of the two haplotypes (Figure 2, Table S2, S3). We also observed fragmented alignments between haplotypes in regions with low gene density and high repetitiveness in several chromosomes (compare panels a-d in Figure 1). Across all 13 chromosomes, SyRI classified only about 71% of the regions between the two haplotypes as syntenic, with around 16% of the genome remaining unaligned (Table S4) due to their extremely repetitive content (see below). While it is not possible to completely exclude that some of these patterns are due to artifacts in sequencing or assembly, these results highlight a high occurrence of structural variation in the stinging nettle genome (Figure 1a, 2b, S2). Structural variation between haplotypes, especially if they have been maintained in nettle for many generations, could also contribute to the high levels of heterozygosity levels we estimated in GenomeScope analyses (5-11%), which are based on short read sequence data from the same stinging nettle individual we sequenced and are therefore estimated independently from the genome assembly. The abundance of SVs in the nettle genome is consistent with the increasingly recognized role of chromosomal structural variants in maintaining sequence and functional diversity in wild [18] and cultivated species [19].

**Figure 2.**
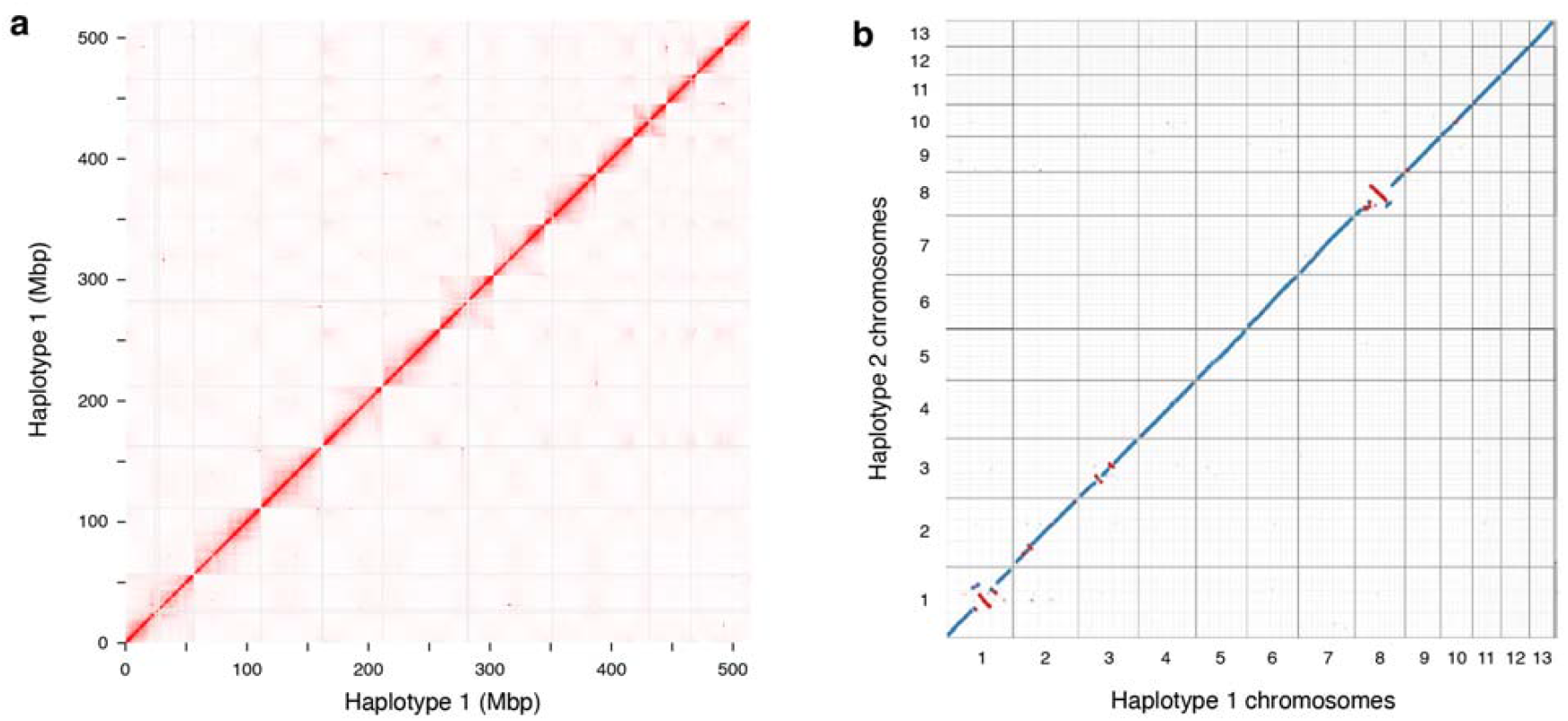
Chromosome structure of the *U. dioica* genome (**a**) Hi-C contact map of haplotype H1 and (**b**) alignment between haplotype H1 and H2 of the *Urtica dioica* ssp. *dioica* genome assembly presented in this study. Only the 13 chromosomes are shown in (**a**); additional smaller contigs were not plotted for clarity. In (**b**), blue represents forward strand alignment, and red represents reverse strand alignment.

**Figure 3.**
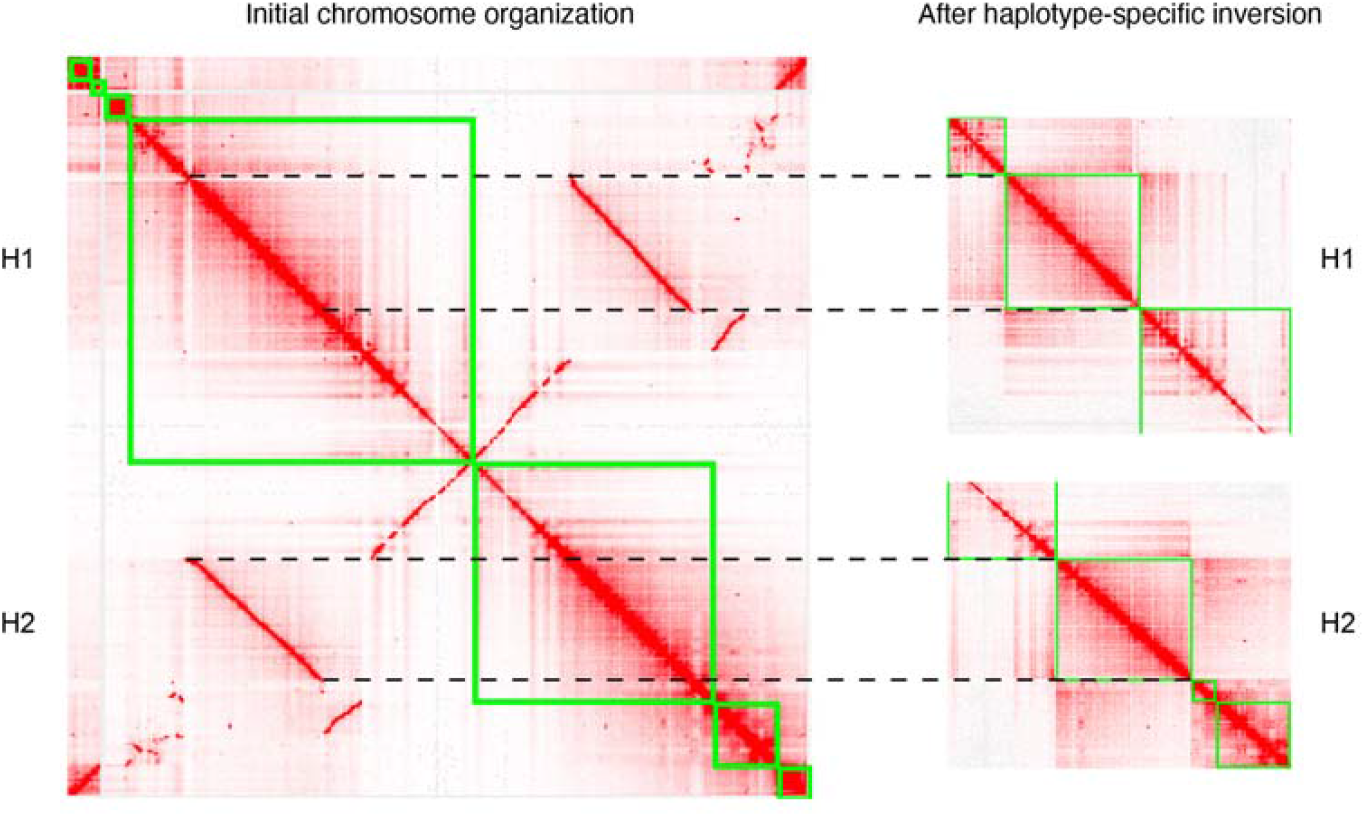
The 18.4 Mbp inversion found between haplotypes on chromosome 8 is supported by Hi-C data. In the left panel, Hi-C data is aligned to both haplotypes simultaneously to create a haplotype-aware heatmap. Only the section of the genome-wide haplotype-aware heatmap corresponding to chromosome 8 is shown; green squares correspond to the H1 (top left) and H2 (bottom right) versions of chromosome 8. The right panel shows changes in the heatmap when Hi-C reads are mapped to modified versions of the assembly in which the orientation of the large putative inversion on chromosome 8 is flipped in H1 (top) or H2 (bottom). Disruption of (haplotype-aware) Hi-C patterns in both of these cases supports the presence of opposite orientations of the inversion in the two haplotypes. Green lines represent contigs.

### 2.3 Genes and repeats landscape

Gene annotation with the BRAKER3 pipeline identified 20,333 and 20,140 genes on H1 and H2, respectively, with annotation BUSCO scores of 90.5% for H1 and 90.4% for H2, showing high levels of completeness (Figure 1b, Table 1, Table S5). Within the 13 chromosomes we identified, we annotated 67.71% and 68.43% of the genome as transposable elements (TEs) in H1 and H2 respectively; we found that the most abundant repeats were Long Terminal Repeat retrotransposons (LTRs), which collectively covered 47.23% and 50.51% of the two assemblies, and Terminal Inverted Repeats (TIRs), which accounted for 16.89% and 13.79% of the assemblies (Table S6). TE density in 500 kbp windows varied from 0 to >6000 TEs per window; however, most windows contained <1000 TEs, except for 37 windows containing more than 2000 TEs (Figure 1c, Table S6).

Scans for patterns of tandem repeats across the genome using RepeatOBserver also identified these TE clusters. RepeatOBserver also calculates a repeats Shannon diversity index, which describes the diversity of tandem repeats across the genome, and uses it to identify the putative location of centromeres in chromosomes. Across most species, centromeric regions have been shown to have high repeat levels but low repeat diversity, and to correspond to minima for the repeats Shannon diversity index (meaning that most of the sequence in these regions is made up of only one or few very abundant repeats). These results can also be confirmed through visual inspection of Fourier transform repeat heatmap, in which discrete banding identifies long stretch of tandem repeats typically associated with centromeres (see, for example, heatmaps for metacentric *Arabidopsis thaliana* (L.) Heynh. and holocentric *Morus notabilis* C.K. Schneid. chromosomes, Figure 4a, b [20–22]). In our *U. dioica* assemblies, repeat patterns are consistent with the presence of acrocentric or near telocentric centromere in five out of 13 chromosomes (chromosomes 8, 9, 11, 12, 13). For eight more chromosomes (chromosomes 1 – 7, 10) we observed more diffused and fragmented patterns of repeats over a large proportion of the chromosome, suggesting the presence of polycentric centromere (Figure 1a, Figure 4c, Figure S4).

**Figure 4.**
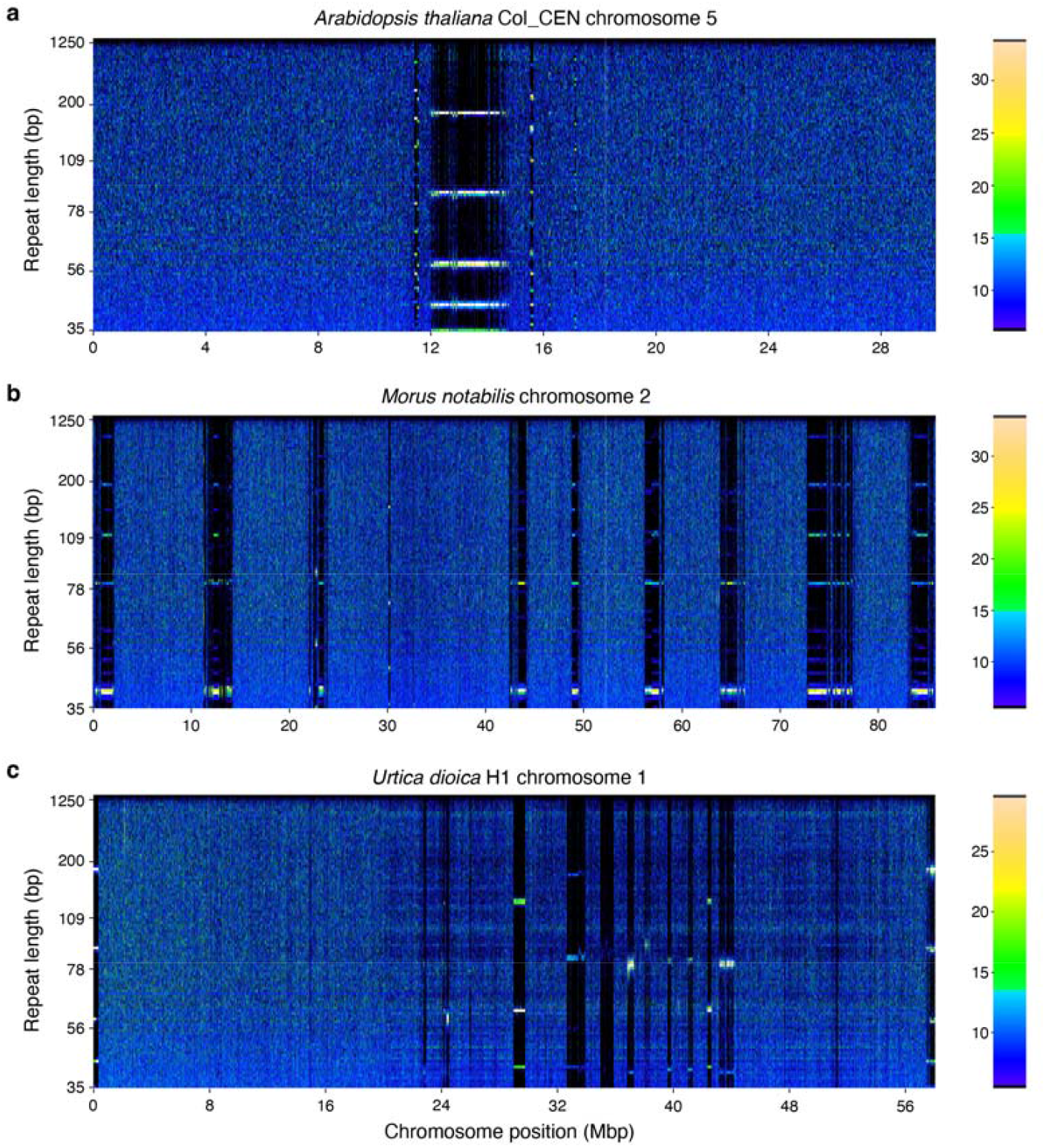
Fourier transform spectra of repeats occurrence in (**a**) *Arabidopsis thaliana* chromosome 5 with metacentric centromeric signal; (**b**) *Morus notabilis* chromosome 2, representative of a holocentric chromosome (Elphinstone et al., 2024); and (**c**) *U. dioica* ssp. *dioica* chromosome 1 (this study). Colour intensity corresponds to the number of times a specific repeat is found in a 5 kbp window. Bright horizontal lines indicate presence of a repeat sequence that repeats itself many times across that region of the chromosome, such as tandem repeats found in telomeric and centromeric regions. Multiple bands in those regions represent harmonics of the base repeat sequence.

While acrocentric centromeres have been previously observed in the genome of another member of the Urticaceae (ramie, *Boehmeria nivea* (L.) Gaudich.; Wang et al., 2021), the presence of polycentric centromeres has not been reported in this family; however, polycentric centromeres have been described in the Moraceae, which are the closest family to the Urticaceae [22]. The position of the predicted centromeres in the *U. dioica* genome is also consistent with those regions having the lowest gene density in the chromosome, as it is typical of centromeres (Figure 1b, e; Table S7). However, it should be noted that these analyses only provide a putative centromere location, and that definitive identification of centromere regions would require direct experimental evidence (e.g. localization of centromeric histones H3, CENH3).

### 2.4 Genome evolution

The haploid chromosome number of *U. dioica* has been variously reported to be 12 or 13 chromosomes [24]. In this study, we unequivocally identify 13 separate chromosomes in the *U. dioica* genome assembly. However, base chromosome number is highly variable within the Urticaceae (ranging from 7 to 14 [25]), as well as within *Urtica* species ([26]. While the Urticaceae family has a complex phylogeny, multiple sources support a monophyletic origin and subdivision in four clades (I-IV [27,28]). The divergence time between clades II/III and clades I/IV was estimated to be 57.43 million years, and the Urticaceae family split from the nearest relative Moraceae 84.87 million years ago [27]. To understand the evolution of genome organization in this family, we compared our *U. dioica* (clade III) genome assembly to those of three other species in the Urticaceae: *Urtica urens* (clade III), *Boehmeria nivea* (clade I), and *Parietaria judaica* (clade I). Despite evidence of abundant large-scale chromosome rearrangements, we found synteny to be quite conserved across all of these species (Figure 5).

**Figure 5.**
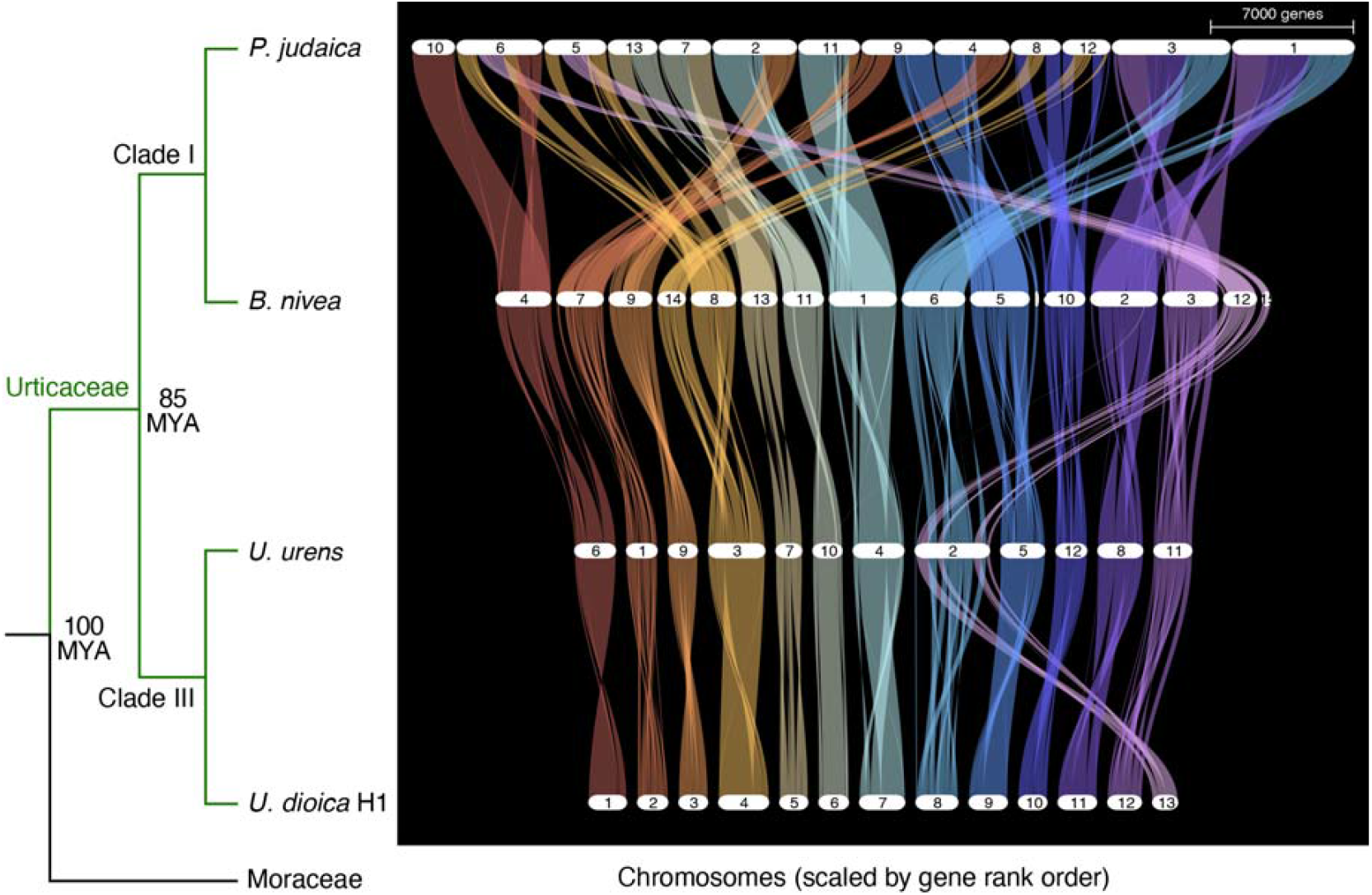
Comparison of chromosome organization in four species of the Urticaceae family. Estimated divergence times are based on Huang et al. (2019). Note that while the *Boehmeria nivea* genome is supposed to have 14 chromosomes, the 15th largest scaffold in the assembly contained more than 300 genes and was therefore included in the figure. While the placement of that section of the genome is quite variable across species, comparison with *Urtica* spp. suggests that it might be part of chromosome 12 in *B. nivea*.

Interestingly, while the phylogenetic distance between *B. nivea* or *P. judaica* and *Urtica* species is comparable, chromosomal synteny is much higher between *Urtica* and *B. nivea* than between either of them and *P. judaica*. Base chromosome number in *U. urens* and many other related *Urtica* species is often n = 12 [25,26], whereas *U. dioica* is almost always n = 13 in diploid cytotypes, while n = 12 is more frequently observed in tetraploids [13]. We find that the *U. dioica* chromosomes 8 and 13 are syntenic with chromosome 2 in *U. urens*, highlighting a clear history of chromosome fusion/fission (Figure 5). The instability of the organization of these chromosomes within *U. dioica* could explain the high level of structural variation that we observe between haplotypes in chromosome 8 of our assembly (Figure 3, Figure S3). As more genomes in this family become available, it will be interesting to further investigate the reasons for this high flexibility in chromosome arrangement between and within related species in the Urticaceae.

### 2.5 Putative neurotoxic nettle sting peptides

Nettle owes its common name to its sting, which constitutes an effective defence mechanism against herbivores. This is due to the presence, on the leaves and stems, of brittle needle-like trichomes, which break upon contact and release pain-inducing chemicals [29]. While the physiological mechanism of the nettle’s sting has been extensively studied since the early 1940s, the pain-inducing compounds were only recently identified. Initial study by Fu et al. [10] showed that simple acids, including oxalic acid, tartaric acid, and formic acid, which are potentially irritant to animals, are the dominant compounds in stinging hairs, suggesting that they could be the pain-inducing compounds in nettle, and rejecting earlier hypotheses [29–31]. More recently, small neurotoxic peptides were shown to play a major role in causing stinging pain. In particular, two classes of such peptides were described in *Urtica* spp. [11]. One is a 42 amino acid-long peptide (4.3-kDa) named urthionin, which has cytolytic activity and a structure that resembles a known group of plant toxins called thionins, known to disrupt the cell membrane [32]. The other is a 63 amino acids-long peptide (6.7-kDa) called urticatoxin, which has neurotoxic activity and is seemingly specific to species in the Urticaceae tribe, such as species in the *Urtica* and *Dendrocnide* genera. Urticatoxin was originally described in *U. ferox* (stinging tree) and was found to induce more severe pain than urthionin. The neurotoxicity of urticatoxin was shown to be due to its ability to modulate the activity of vertebrate ion-gated sodium channels, similar to the gympietides (i.e., Excelsatoxin and Moroidotoxin) previously described in *Dendrocnide* species [12]. Despite their similar effect, urticatoxins and gympietides appear to have evolved independently, given their structural differences [11].

Using a homology-based approach, we identified two copies of a gene putatively encoding the urthionin peptide on chromosome 9 (86% amino acid match to the *U. ferox* mature peptide: •-Uf1a). The two paralogs were positioned right next to each other with the same sequences, indicating a recent gene duplication event. Hits for urticatoxin on chromosomes 6 and 9 were only a 26-49% match to the peptides identified in *U. ferox* and in *Dendrocnide* species (Table 2), suggesting that this class of peptides might not be found in *U. dioica*. No genes with similarity to two other neurotoxic peptides identified in *Dendrocnide* species

**Table 2:**
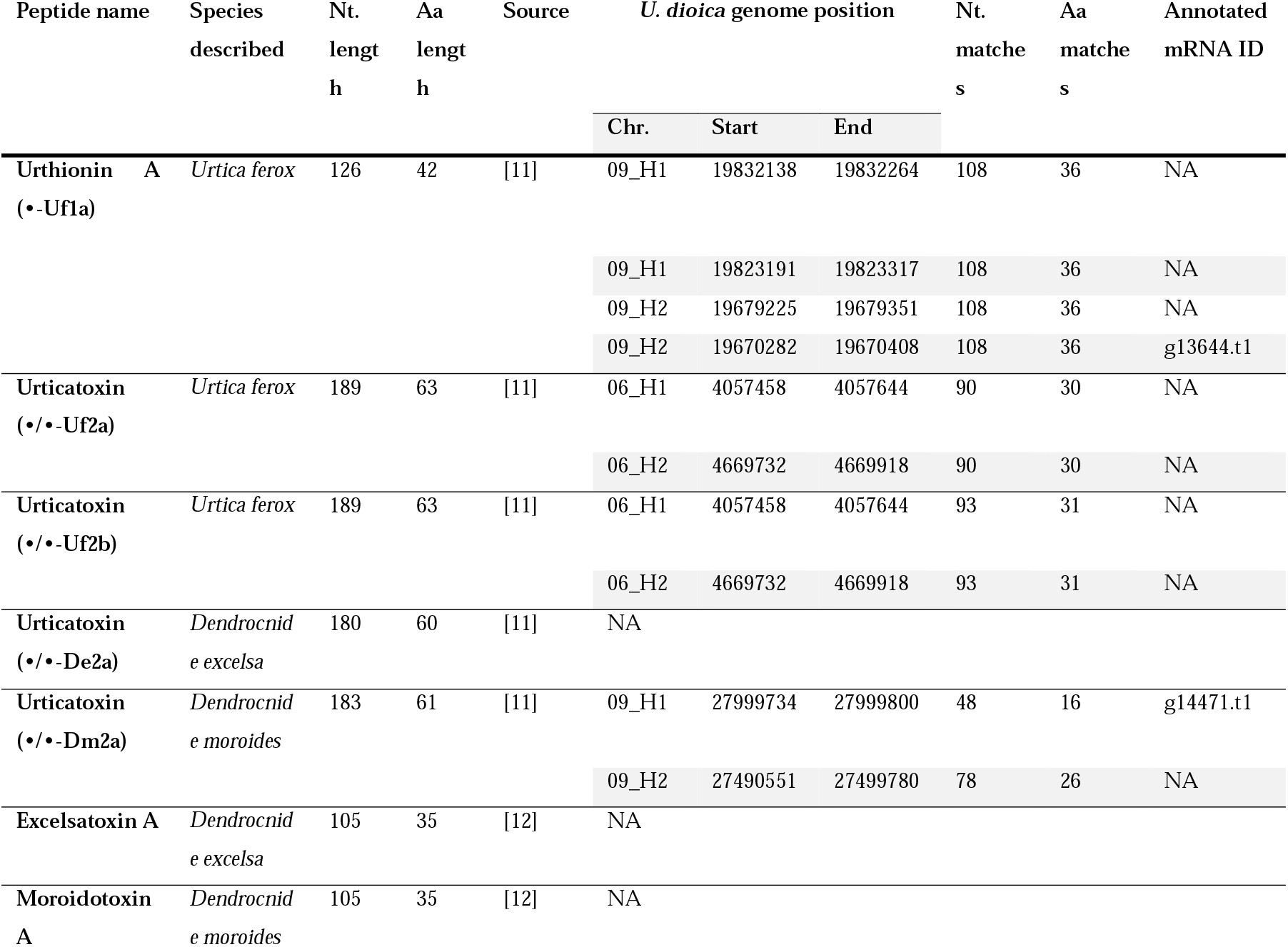
Identification of *U. dioica* homologs of pain-inducing peptides identified in other stinging species. Nt. = nucleotides; Aa = amino acids.

(Excelsatoxin A and Moroidotoxin A) were identified in the nettle genome.

## 3. Materials and Methods

### 3.1 Plant materials and sequencing

We collected a diploid female individual of *Urtica dioica* ssp. *dioica* from beside the River Jiu, north of Rovinari, Romania, and cultivated it in Vancouver, British Columbia (as clone 11-4). Young leaves were collected and flash-frozen in liquid nitrogen for extractions. A voucher specimen was deposited in the herbarium of the Beaty Biodiversity Museum at the University of British Columbia (UBC).

Estimation of nuclear DNA content was achieved using an Attune NxT Flow Cytometer (Thermo Fisher Scientific) at the University of British Columbia; preparation of samples followed the protocol in [33] using a Tris-MgCl_2_ lysis buffer [34]. RNA was removed with RNase A prior to staining with propidium iodide [33], and a tomato (*Solanum lycopersicum* L.) sample prepared with the same method was used as a standard when estimating the nettle genome size.

High-molecular-weight DNA was extracted using a modified CTAB method [35]. A PacBio HiFi sequencing library was prepared and sequenced on a Revio instrument by Novogene (San Diego, CA, USA). HiFi reads were quality-controlled using SMRT tools v13.0.0, runqc-reports (PacBio, 2024). Whole genome shotgun sequencing (WGS) was performed on an Illumina HiSeq 2000 platform by CD Genomics (Shirley, NY, USA) to produce 150 bp paired-end reads. DNA for Nanopore sequence was extracted using a ThermoFisher MagMax Plant DNA kit, (Waltham, MA, USA) and then further purified using a Qiagen DNeasy PowerClean column (Hilden, Germany). Nanopore sequencing libraries were generated from HMW DNA using the Genomic DNA by Ligation (SQK-LSK110) protocol. Sequencing was carried out with FLO-FLG001 R9.4.1 flow cells on a MinION instrument. The resulting fast5 files were subsequently basecalled using Guppy 6.0.1 Superior Basecalling Algorithm (dna_r9.4.1_450bps_sup.cfg) conducted on an NVIDIA 3060ti. Reads with a Q score below 9 were discarded.

A Hi-C library was prepared with modifications. In brief, ground frozen tissue was cross-linked in a 1.5% formaldehyde solution containing protease inhibitor. Following nuclei isolation, chromatin was fragmented through DpnII digestion. After end-filling in the presence of biotinylated dATP, blunt ends were ligated, and DNA was extracted. Three and a half μg of DNA were sheared using ultrasonication (Covaris, Woburn, MA, USA), and fragments in the 300-500 bp size range were further selected using SPRI beads [36]. Biotinylated fragments were pulled down using streptavidin-coated beads (Invitrogen, Waltham, MA, USA), and Illumina libraries were prepared following Todesco et al. [37]. The Hi-C method used here combined elements from various previously published Hi-C protocols (including [38,39]) and has been optimized to work on a variety of plant species and to reduce library preparation costs. A detailed protocol is provided in the Supplementary Material. The resulting libraries were sequenced by Novogene (San Diego, CA, USA) on an Illumina NovaSeq X Plus instrument to generate 150 bp paired-end reads.

### 3.2 De novo assembly and quality evaluation

We tested multiple assemblers, assembly parameters and versions to determine what worked best for our nettle assembly, and selected the approach that produced the most complete kmer representations in both haplotypes, based on a kmer plot analysis with Merqury [40]. PacBio HiFi reads were first filtered by mean Q score >20 using fastq.filter -e 0.01 (https://github.com/LUMC/fastq-filter). An initial contig-level genome assembly was produced with hifiasm v0.19.8-r-603, integrating Hi-C reads using the --h1 and --h2 options [41] and keeping default values for the remaining parameters. This resulted in two separate haplotype assemblies. Juicer [42] and 3D-DNA [43] were used to map Hi-C reads and create a contact map, which was used to manually sort and orient the contigs to produce a chromosome-scale assembly for each haplotype in Juicebox [44]. To verify the positions and orientation of contigs, we aligned the two haplotypes and compared syntenic regions. Alignments were done using minimap2 v2.28 [45,46] and regions harbouring putative structural variants (SVs) were surveyed for obvious misassemblies on the Hi-C contact map. For regions showing ambiguous patterns in the Hi-C contact alone, we used Synteny Rearrangement Identifier (SyRI) v1.7.0 [47] to obtain precise coordinates of the putative SVs. SVs larger than 10,000 bp were then manually reoriented in individual haplotypes to determine if that would improve the Hi-C contact profiles in the region (Table S2). SVs smaller than 10,000 bp could not be visually assessed on Juicebox and were therefore omitted from this manual curation step. To check whether these putative SVs were supported by the presence of long reads spanning the expected breakpoints (following Harringmeyer & Hoekstra, 2022), we then mapped filtered HiFi reads and ONT reads •30 kbp to the assembled genome using winnowmap v2.03 [49] and visualized the resulting alignments on IGV (Table S3). We used this evidence to curate both haplotype assemblies and correct likely misassemblies. Furthermore, we re-mapped the Hi-C reads to produce a haplotype-aware H1+H2 contact map using 3D-DNA with parameter -q 0 [43] to finalize the assembly, which allowed for visualization of reads that were mapped to multiple regions in the genome (i.e., mapping quality of 0). This allowed to resolve highly repetitive regions that previously showed no visible interactions in Juicebox. However, we note that this final step fragmented our genomes into smaller contigs as a trade-off for smoother Hi-C contact map patterns. We then aligned our assemblies to the primary haplotype of a published *U. dioica* assembly (NCBI accession: GCA_964188135.1, hereafter presented as Udio_DToL) to check for synteny, and we assigned corresponding chromosome numbers. To verify whether differences in chromosome organization with respect to Udio_DToL could reflect misassemblies in our haplotypes, we re-ordered our contigs based on Udio_DToL using Ragtag v2.1.0 [50] (command: ragtag.py correct + ragtag.py scaffold), re-mapped our Hi-C reads and generated a new contact map on Juicebox. If our assembly is a better representation of the real order and orientation of the contigs for our sequenced individual, regions that have a different organization in this reference-based scaffolding assembly should appear as misassemblies in the Hi-C contact map (Figure S3).

Genome assembly statistics were assessed with Bandage v0.8.1 [51], BBmap v39.06 [52], and BUSCO v5.1.2 (dataset: eudicot_odb10 [53,54]). Additionally, we used the Illumina WGS data to calculate kmer completeness and QV score with Merqury [40]. In brief, we removed adapters and kept only the paired reads from the raw Illumina data using Trimmomatic v0.39 with parameters ILLUMINACLIP: TruSeq3-PE.fa:2:30:10:2:True SLIDINGWINDOW:4:15 LEADING:3 TRAILING:3 MINLEN:36 [55]. Then, the meryl database was built using the trimmed Illumina reads with k = 19, based on which Merqury generated the kmer plots for quality assessment (Figure S1). Heterozygosity was calculated using GenomeScope2 with k = 21 [56]. We observed a significant portion of the contigs unplaced to the chromosomes, lacking any interactions with chromosomes but interacting strongly with themselves on Hi-C contact map. To assess whether these unplaced contigs contain genes or are mostly composed of repeats, we ran the same BUSCO analysis as above and the redmask.py v0.0.2 command in Red with default parameters to identify repetitive regions [57].

### 3.3 Genome annotation and visualization

We performed annotation of our stinging nettle genome assembly using BRAKER3, which allows integration of RNAseq data to support the annotation process and has shown superior benchmarking performance in published studies [58–60]. To run the pipeline, we first soft-masked the repetitive regions in the genome using the redmask.py v0.0.2 command in Red, with default parameters [57]. RNAseq data from three different tissue types - leaf, fibre, callus - was retrieved from Xu et al. [61]; raw paired-end Illumina reads were filtered with Trimmomatic v0.39 with parameters ILLUMINACLIP: TruSeq3-PE.fa:2:30:10:2:True SLIDINGWINDOW:4:15 LEADING:3 TRAILING:3 MINLEN:36 [55]. Filtered read pairs were aligned to the soft-masked genome using Hisat2 v2.2.1 (parameters --dta added), and the alignment file was converted to BAM format using samtools [62]. Prior to moving forward, the alignment score of the RNAseq to the genome was checked to be above 80% on average to confirm that the individuals from which the RNAseq was obtained from were sufficiently similar to our *U. dioica* assembly. The resulting BAM file, as well as the Viridiplantae protein database from OrthoDB v11 [63,64], were incorporated in the BRAKER3 pipeline (using --bam and --prot_seq mode). Completeness of the annotation was assessed with BUSCO with --mode protein (dataset: eudicot_odb10 [53,54]).

For transposable elements (TE) annotation, we used the extensive *de-novo* TE annotator (EDTA) pipeline v.2.2.0 [65] with default parameters. Additionally, we used RepeatOBserver v1 [20] to analyze patterns of tandem repeats across the chromosomes and identify putative telomeric and centromeric repeats. The location of the putative centromere was predicted in RepeatOBserver using the default values for Shannon diversity standard deviation cut-off (Shannon_bin_size=500, Shannon_SD=2). Genome-wide patterns of gene and TE density, Shannon diversity and rearrangements between haplotypes, as well as putative centromere locations, were visualized using Circos v0.69-8 [66]; computed Shannon diversity scores were averaged over a 250 kbp window into a line plot, and the gene and TE annotations were formatted to histograms of counts per 500 kbp window. To ensure proper visualization of TE patterns across the genome, we set the maximum y-axis for the TE density track to 4000, resulting in the values for six outlier windows with very high TE counts (>4000) being cut off the plot. Location and TE counts in those regions is reported in Table S6.

### 3.4 Comparative genomics

To investigate patterns of chromosome evolution within the nettle family, we first compiled a list of reference genomes published within the Urticaceae. The quality of these genome assemblies was assessed based on the annotation BUSCO score (•90%). We included *Urtica urens* L., *Parietaria judaica* L. (both primary haplotypes from the Darwin Tree of Life Project Consortium [17]), and a wild accession of *Boehmeria nivea* [23]. For each genome, the gene annotation file in gff format was downloaded together with the associated protein sequences in amino acid fasta format. Then, the annotation was converted into a bed format using the lines corresponding to the “mRNA” annotation, with an extra column matching the header of the protein fasta sequence file. This bed file then indicates the genome position of every protein. File conversion was done manually with a custom shell script as needed. For this analysis, we used the H1 haplotype of our *U. dioica* assembly since it had a longer total assembly size (quality scores are otherwise similar between the two haplotypes). Once the input was prepared for all the downloaded datasets and our H1 genome assembly, we ran GENESPACE v1.3.1 [67] to compare chromosome synteny across the four Urticaceae species. GENESPACE was run with default parameters. In brief, initial orthogroups were discovered and identified with OrthoFinder v2.5.4 [68]. Using the synteny information analyzed with MCScanX [69], syntenic blocks based on the gene orders were visualized.

### 3.5 Identification of sting genes

To identify putative neurotoxic peptides in our stinging nettle genome, we obtained the amino acid sequence of the 4.3-kDa urthionin (•-Uf1a) and several versions of 6.7-kDa neurotoxic urticatoxin (•/•-Uf2a, •/•-Uf2b, •/•-Dm2a, •/•-De2a) peptide, described in *Urtica ferox* G. Forst, *Dendrocnide moroides* (Wedd.) Chew and *D. excelsa* (Wedd.) Chew [11]. Xie et al. described the above toxins in other *Urtica* species, including from transcriptome data of *U. dioica* and *U. incisa* Poir.; however, since these putative toxin transcripts were found by homology to *U. ferox* sequences, we omitted them from our analysis. Additionally, we included in our search the two 4-kDa gympietides (Moroidotoxin A, Excelsatoxin A) described in *D. moroides* and *D. excelsa* [12]. After compiling this list of seven sting-associated neurotoxic peptides found in the literature, we looked for possible homologs in stinging nettle by aligning their protein and transcript sequences to our genome assembly using miniprot v0.13 (r248) with the parameter -Iut16, which accounted for a flexible size range of introns in the homolog detection on the query genome [70]. The visualized alignment results of these analyses are reported in Figure S5.

## 4. Conclusions

We have generated a highly complete, diploid genome assembly of stinging nettle, a multi-purpose species whose use has been interwoven in various cultures for thousands of years and that is seeing renewed interest as a source of natural, sustainable fibre. Despite its compact size, we found the stinging nettle genome to be highly repetitive, with almost two-thirds of its sequence being composed of transposable elements. We also identify surprisingly high levels of structural variation between haplotypes in the individual we sequenced; while, despite our best efforts, we cannot completely exclude that these are due to haplotype misassemblies, it seems likely that these two observations are linked, given that transposable elements are known to facilitate the generation of chromosomal rearrangements [71]. Further complexity in the nettle genome is added by the possible presence of several polycentric chromosomes, as suggested by diffuse patterns of short tandem repeats. The stinging nettle genome represents, therefore, an attractive case study to help understand the evolution of chromosomal variation within and between species; it will be interesting, in future studies, to determine if the inversions we identified play a role in ecotypic adaptation in nettle, as has been observed in other plant species [72]. This assembly also provides a valuable resource for the nettle community, as it will greatly facilitate analyses of the genetic makeup of useful and interesting traits in the nettle family, such as studies of the genes controlling fibre quality, the production of bioactive molecules found in this species, or the formation of nettle’s characteristic stinging hairs.

## Supporting information

Hi-C protocol

## Author Contributions

Conceptualization: QC and MD; funding and resources: QC, MD, GG, DP and MT; data production: KH, CD, MK, DP; formal analyses, investigation, and visualization, KH and MT; sample preparation and laboratory work: KH, CD, MK, DP; writing, review and editing: all authors. All the authors read and approved the final manuscript.

## Data Availability Statement

The raw sequencing data and the assembled genome are available at NCBI under BioProject number PRJNA663211. All the scripts used in this study are available at https://github.com/kaede0e/stinging_nettle_genome_assembly.git. Clones of individual 11-4 are available upon request.

## Acknowledgments

We thank Eric Gonzales Segovia and Cassandra Elphinstone for help with genome scaffolding and tandem repeats analyses, respectively, and Andy Johnson and Cassandra Elphinstone for assistance with the flow cytometry. This study was supported by NSERC Discovery Grants to MT (RGPIN-2023-03344), QC (RGPIN-2019-04041), and MD (RGPIN-2020-05147). We also gratefully acknowledge the Digital Research Alliance of Canada (DRAC) for access to their computational resources, and the Natural History Museum, London (UK) for funding the fieldwork in Europe undertaken by DP and QC.

