## Supplemental Figures for "A high-quality phased genome assembly of stinging nettle, *Urtica dioica* ssp. *dioica*"

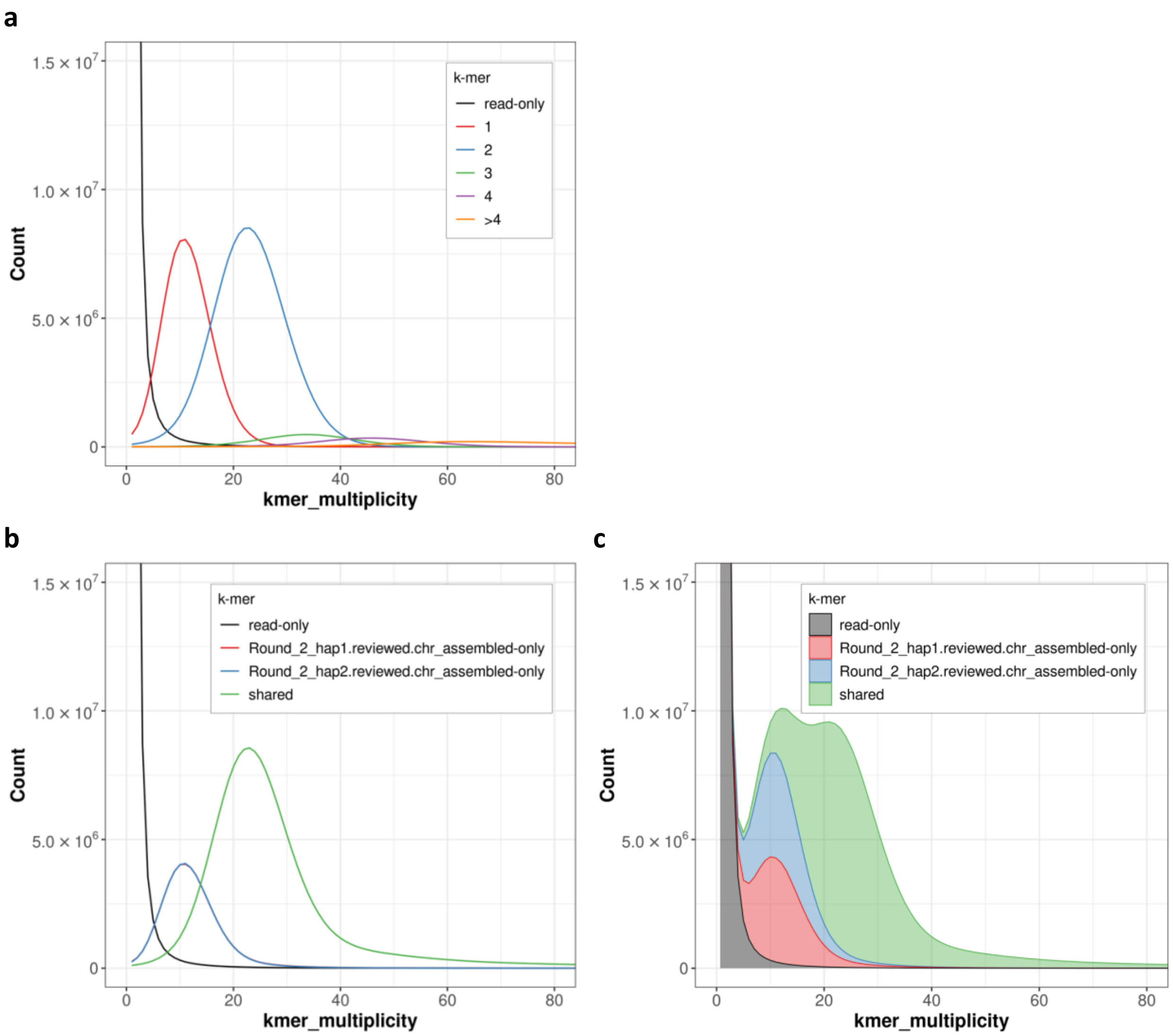


**Figure S1:** Kmer plot for the *U. dioica* ssp. *dioica* assembly produced with Merqury using WGS Illumina data. (**a**) Unstacked histograms of kmer counts per copy numbers found in the assembly. The first peak corresponds to the kmers found only once in the genome (haploid, meaning that they correspond to heterozygous region) and the second peak corresponds to the diploid peak (homozygous regions). (**b**) Kmer counts sorted by haplotype assembly. The line corresponding to H1 (red) line is barely visible in the plot because it almost completely overlaps with the H2 line (blue; first peak in the plot). This indicates that both haplotype assemblies are equally complete in their kmer presence. (**c**) Stacked histogram for **b**.

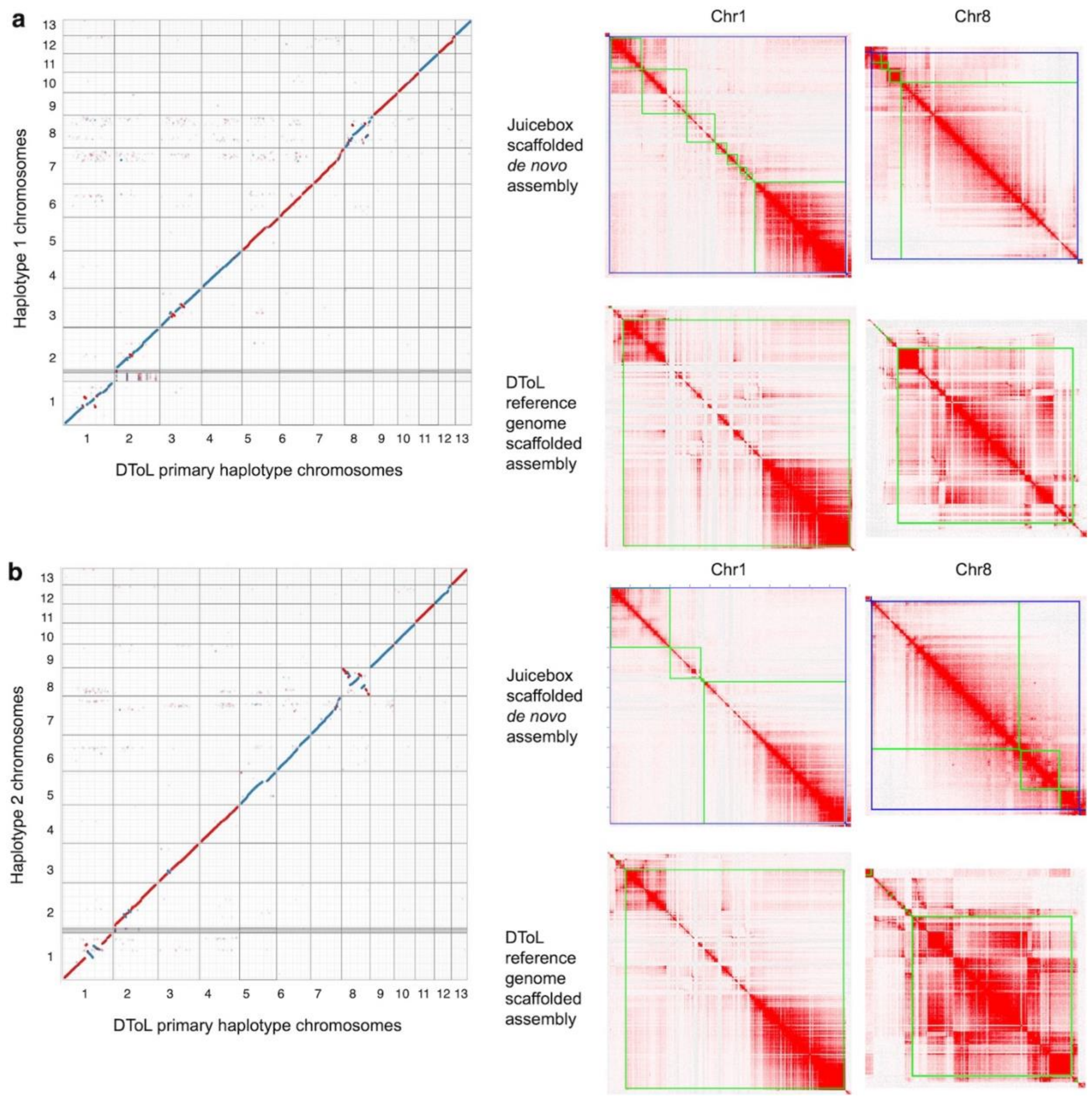


**Figure S2:** (**a**) Alignment between the *U. dioica* assembly from Darwin Tree of Life (DToL) and the *U. dioica* ssp. *dioica* H1 and b) H2 assembly from this study, done using minimap2 (left panel). Given the numerous structural differences between our study and DToL assembly, we wanted to assess whether the DToL assembly was a better overall representation of the genome organization in *U. dioica*. To do so, we re-scaffolded the contigs in our assembly using the DToL *U. dioica* genome primary haplotype assembly as reference, using RagTag (correct + scaffold). We then mapped our Hi-C data to the re-scaffolded assembly. In all cases, re-scaffolding using the DToL genome assembly as reference resulted in less clean contact maps as seen in the two example chromosomes shown on the right panels. This validates the chromosome organization of our assembly, and suggests the presence of several major structural variants between the genome of the *U. dioica* individual we sequenced and the one that has been sequenced by DToL.

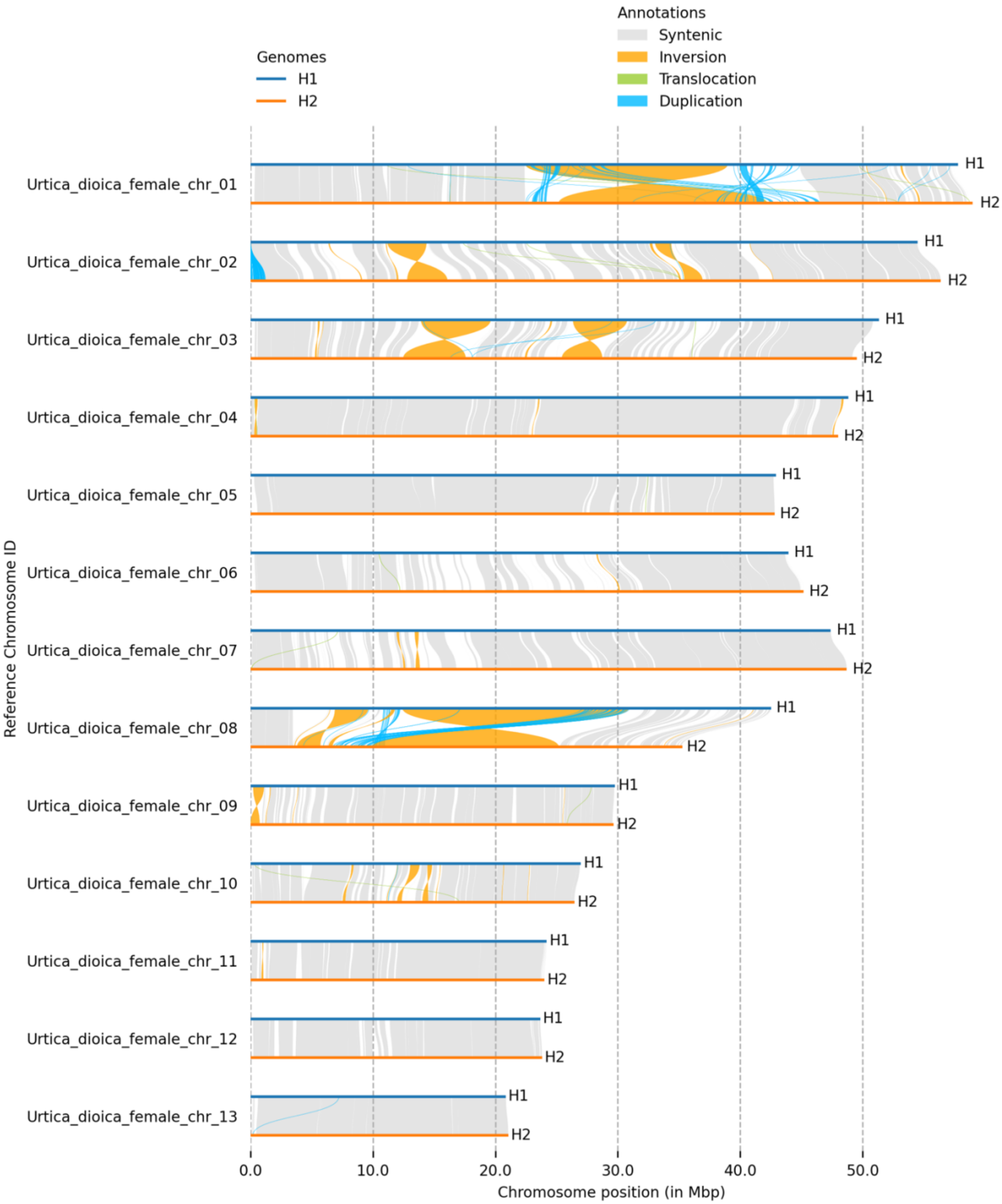


**Figure S3:** SyRI alignment between *U. dioica* ssp. *dioica* H1 and H2 haplotypes. The plot was generated with plotsr.

## H1

Chr1

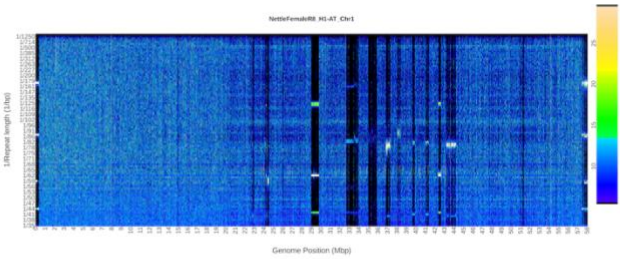


Chr2

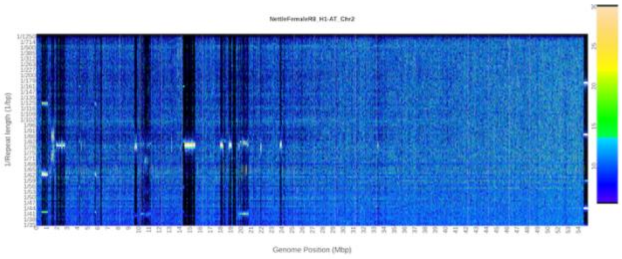


Chr3

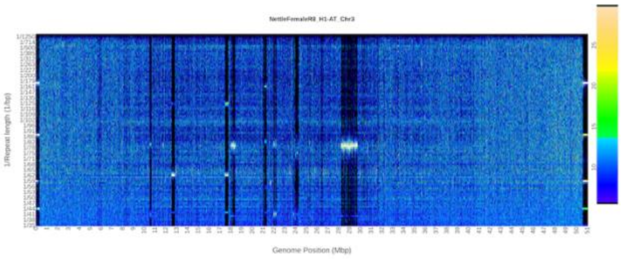


Chr4

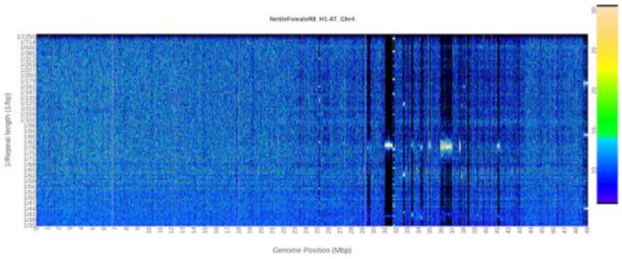


Chr5

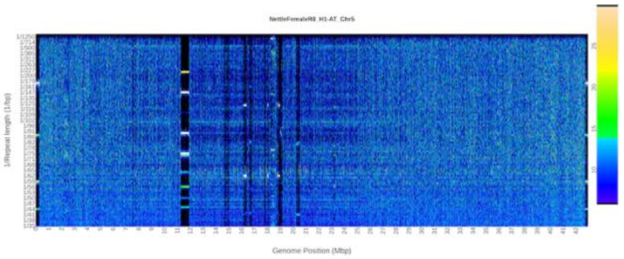


Chr6

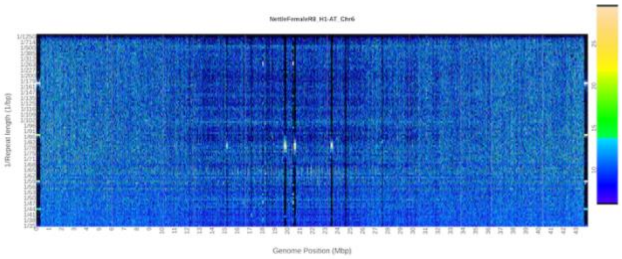


Chr7

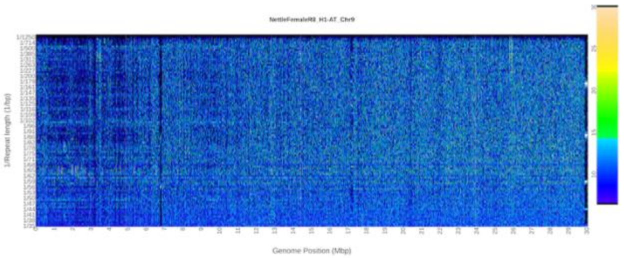


Chr8

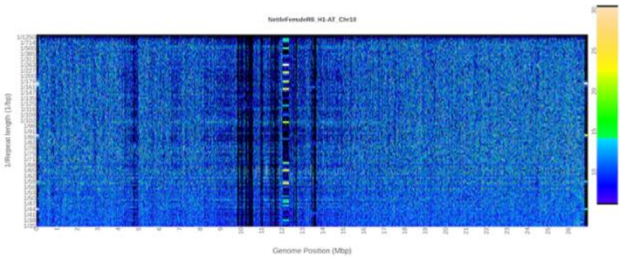


Chr9

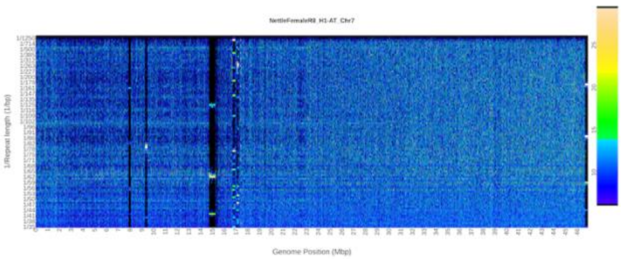


Chr10

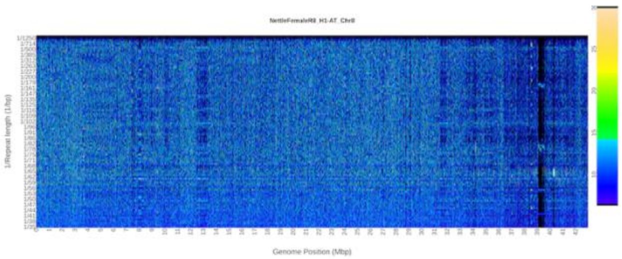

Chr11

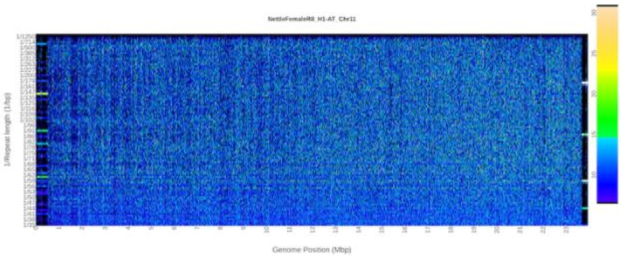


Chr12

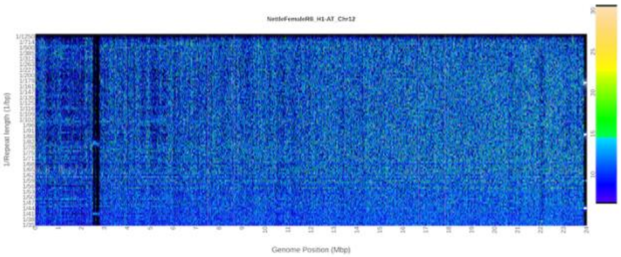


Chr13

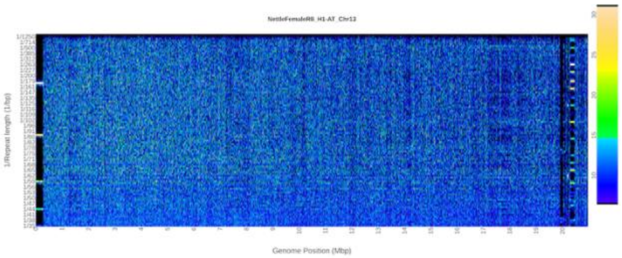


## H2

Chr1

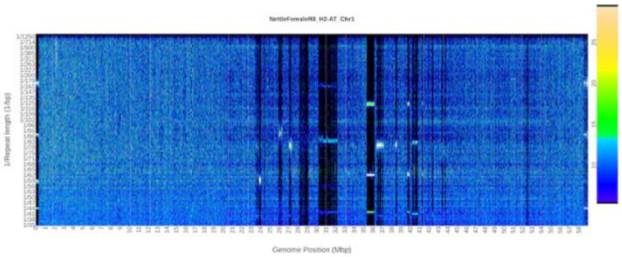


Chr2

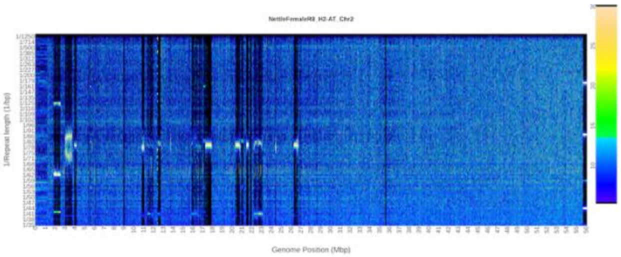


Chr3

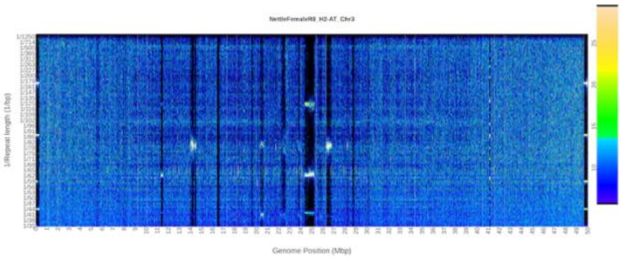


Chr4

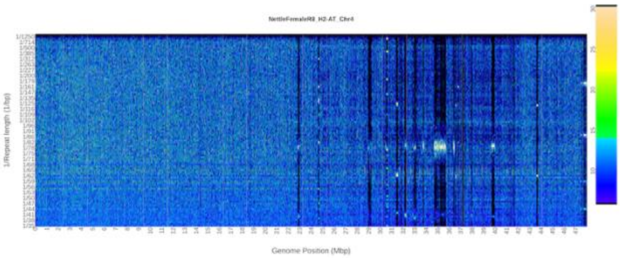


Chr5

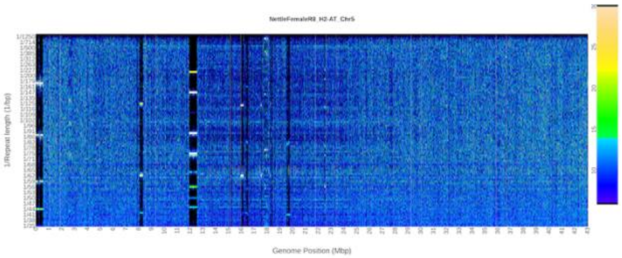


Chr6

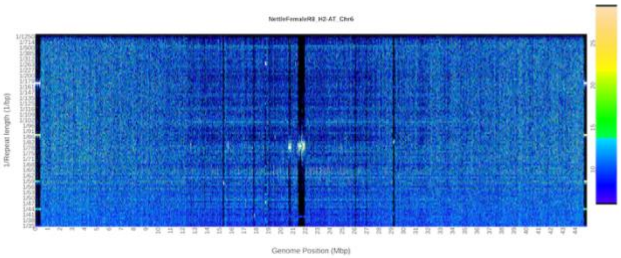

Chr7

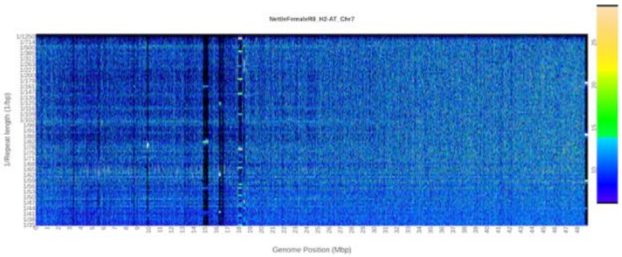

Chr8

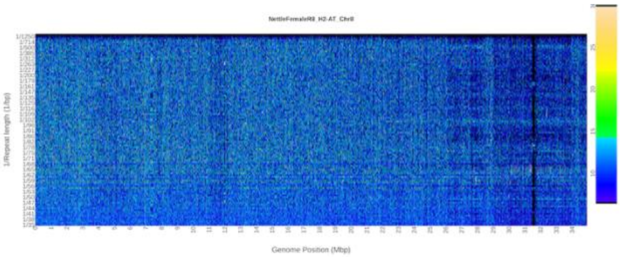

Chr9

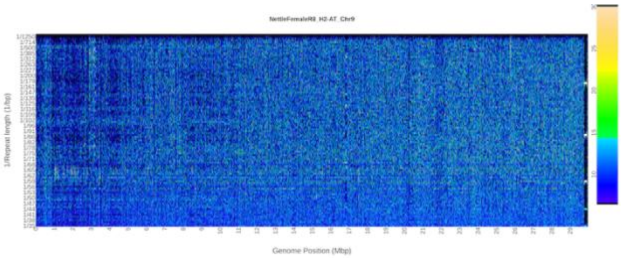

Chr10

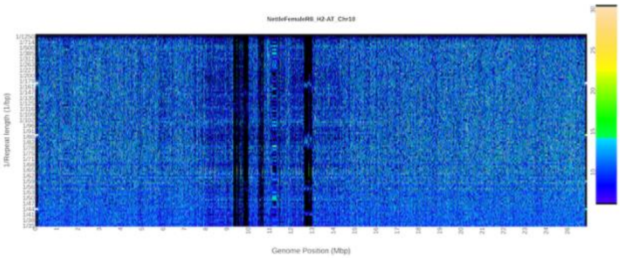

Chr11

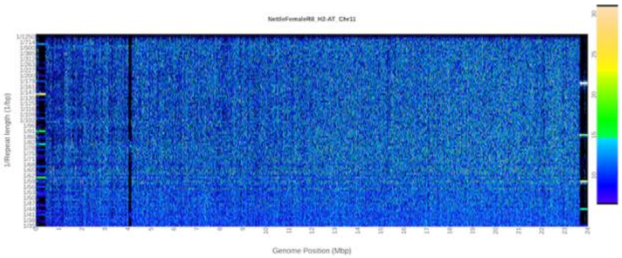

Chr12

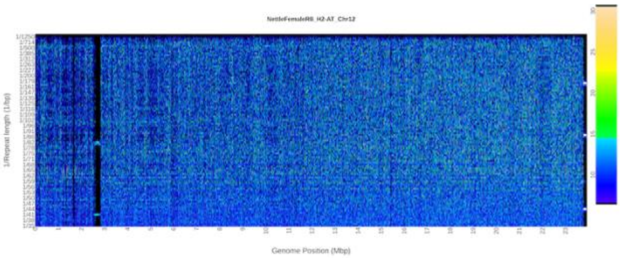

Chr13

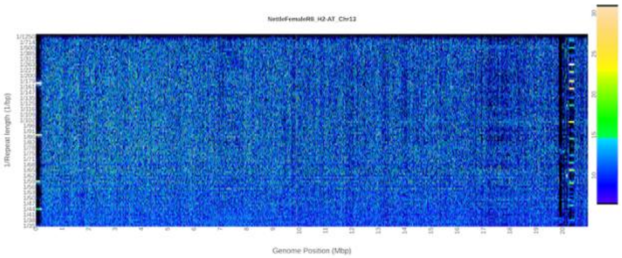

**Figure S4:** Fourier transformed repeat spectra for each chromosome obtained with RepeatOBserver. The x-axis represents the position on the chromosome, and y-axis represents 1/repeat length. Colour intensity corresponds to the number of times a specific repeat is found in a 5 kbp window. Regions with high proportions of tandem repeat show characteristic banding patterns (bands corresponding to larger repeats sizes are harmonics of the smaller base repeat).

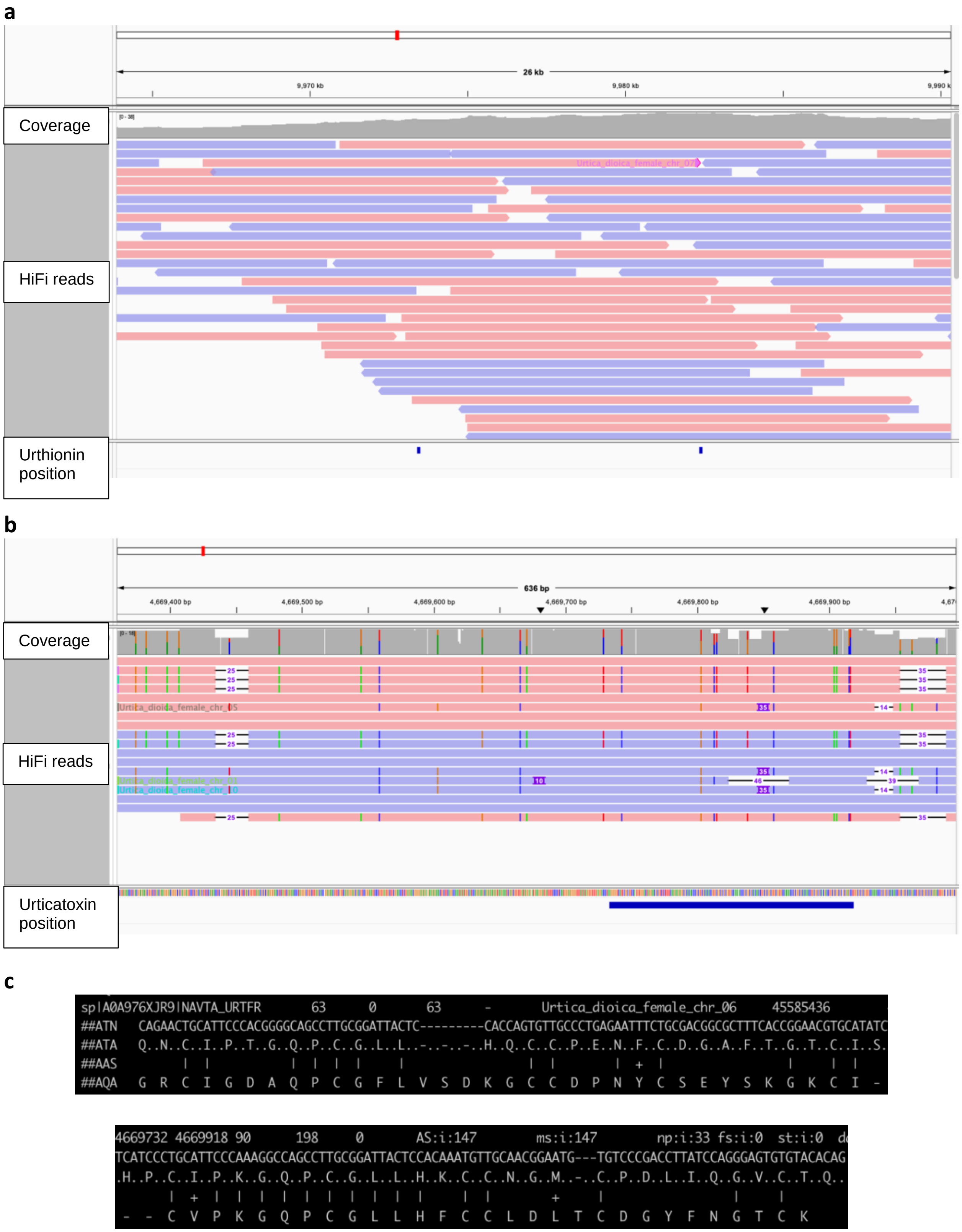


**Figure S5:** Confirmation of the putative stinging peptide genes. Positions of the putative (**a**) urthionin and (**b**) urticatoxin genes on the *U. dioica* genome, visualized in IGV. Alignment of HiFi reads (middle track) against the genome assembly indicates that these are well-assembled region with good coverage (top track). Several long reads support the presence of the putative stinging peptide genes in these regions (bottom track). (**c**) Amino acid alignment between urticatoxin from *U. ferox* (Xie et al. 2022) and the putative urticatoxin peptide in the *U. dioica* genome.
