## Supplementary material for "A high-quality phased genome assembly of stinging nettle, *Urtica dioica* ssp. *dioica*": Hi-C protocol

### Plant Hi-C Library Preparation

Adapted and optimized from Padmarasu *et al.*[1] and Wang *et al.*[2]

This protocol has been successfully tested in more than a dozen different species; besides nettle, cultivated sunflower and several of its wild relatives, various *Rubus* and *Vaccinium* species, cannabis, Douglas fir, *Amaranthus tuberculatus*, among others, as well as on kelp species. The efficiency of nuclei isolation can be quite variable depending on the species, and we recommend testing that (see note on step A.1) before committing to the full protocol. The nuclei isolation buffer we report here worked well for all of the species above, but a different buffer might be required for more difficult-to-handle species. For example, we found that further buffering the nuclei isolation solution by increasing the concentration of Tris can help with nuclei and DNA recovery from some long-lived arctic species. It should also be noted that the nuclei obtained in part B are not particularly clean and might contain quite a bit of cell debris, making it often difficult to visualize them clearly on a microscope; the scope of that step is to isolate intact nuclei from which DNA will be then extracted, and cell wall debris do not interfere with the process.

The protocol is fairly lengthy, because it includes a few overnight incubation, but several of the days require little input in terms of time. While it is likely possible to shorten those incubation times with minimal impacts on the final results, we have not thoroughly optimized that.

At various steps during the initial part of the protocol it is necessary to keep some of the material as controls, to make sure nuclei isolation, digestion and ligation in the nuclei worked properly. Do not skip those controls!

This protocol was developed by Marco Todesco and further optimized by Natalia Bercovich and Kaede Hirabayashi.

#### A. Fixation and nuclei extraction

##### Tissue collection and grinding (can be done ahead of time)

1. Collect enough tissue to yield 2-10 ug of DNA after nuclei fixation and extraction. Very young, developing (possibly etiolated) leaves are preferred. For most species we have used this protocol on, 0.35 to 1 g of very young leaves has been sufficient. For mature leaves or other tissue, more starting material might be needed, as is the case for more problematic species (for kelp samples, we use 5 g of mature tissue).

   **Note:** It is a good idea, if extra tissue is available for the individual of interest or for another individual of the same species, to perform a preliminary tests to assess efficiency of nuclei extraction and DNA isolation. To do this, perform steps A.1 to B.5, and the extract DNA from the isolated nuclei following steps F.1 until F.16 (ignoring the CONTROL step), and then measure concentration using a Qubit HR kit.

2. Flash freeze in liquid $N_2$ and grind thoroughly using pre-chilled mortar and pestles. Once the tissue if homogenously pulverized, add to the mortar ~25 ml of liquid $N_2$ (if the pulverized tissue is stuck to the mortar, scrape it off with a pre-chilled metal spatula beforehand). Pour the pulverized tissue, resuspended in liquid $N_2$, into a 50 ml conical tube. Use a pre-chilled metal spatula to remove any remaining tissue from the mortar and add it to the 50 ml conical. Close the lid loosely, and store in a -80 ºC freezer until the $N_2$ evaporates. Close the lid tightly, and proceed with the protocol or store at -80 ºC until needed.

#### Hi-C DNA preparation

##### DAY 1

3. For each sample, **<u>prepare ahead</u>** 20 ml of NIB and supplement it with:

   500 µl of 10% Triton X-100 (0.25% v/v final)

   200 µl of 50 mM spermine (0.5 mM final)

   20 µl 100 mM PMSF (0.1 mM final)

   20 µl beta-mercaptoethanol (0.1% v/v final)

   200 µl protease inhibitor cocktail (1% v/v final)

4. Add 15 ml of pre-cooled 1x PBS plus 607 µl of 37% formaldehyde (FA) into the tube containing the plant powder (1.5% v/v FA final concentration). The 37% FA solution should not have been opened for more than three months.
5. Vortex mix and incubate at room temperature (RT) for 15 minutes, with gentle rotation.
6. Add 2 ml of glycine 2M (0.250 M final concentration), incubate at RT for 5 minutes with gentle rotation. Put tube on ice for 5-10 minutes.
7. Pellet sample at 2,000 × g for 5 minutes at 4 ºC.
8. Wash once with cold 1x PBS (20 ml), pellet at 2,000 × g for 5 minutes.
9. Remove supernatant, add 10 ml of cold Nuclei Isolation Buffer to which the solution mentioned in A.3 have been added (NIB+).
10. In a cold room, filter through two layers of cheese cloth pre-wetted with NIB+, and then one layer of miracloth pre-wetted with NIB+.
11. Spin at 3,500 × g for 10 minutes at 4 ºC.
12. Resuspend in 1 ml of NIB+, <u>gently</u> agitating with a pipette tip to resuspend the pellet.
13. Move to a 2 ml tube using a wide-bore pipette tip (on ice).
14. Centrifuge at 2,000 × g for 5 minutes at 4 ºC.
15. Discard the supernatant. If there is a green slurry on top of the grey pellet of nuclei/fibers/starch, remove with a pipette.
16. Resuspend gently in 1.5 ml of NIB+.
17. Spin down at 2,000 × g at 4 ºC for 5 minutes and then remove the supernatant by pipetting, making sure not to disturb the pellet.
18. Repeat wash with 1.5 ml of NIB+ (steps A.16-A.17).

##### B. Nuclei fractionation

1. Resuspend (gently) in 2 ml of cold Percoll gradient buffer (Add 1 ml of the Percoll gradient buffer; if needed, gently dislodge the pellet from the bottom of the tube by mixing with a pipette tip, and mix by gentle inversion. Add another 1 ml of the Percoll gradient buffer, and mix again by gentle inversion).
2. Spin at 12,000 × g for 15 minutes at 4 ºC.
3. Scoop the nuclei floating on the surface of the Percoll gradient with a clean spatula and put them in a 2 ml tube containing 1.5 ml of cold NIB+. Collect what is left of the nuclei on top of the Percoll gradient with a pipette (wide-bore tips) and add it to the same 1.5 ml tube. Resuspend pellet the nuclei pellet by <u>gently</u> swirling with pipette tip.
4. Spin at 2,000 × g for 5 minutes at 4 ºC. Discard the supernatant **CAREFULLY** – the pellet might not be solid due to remaining Percoll. If so, remove some of the supernatant with a pipette, add 1 ml cold NIB+, and mix by gentle inversion. Spin again at 2,000 × g for 5 minutes at 4 ºC and remove the supernatant.
5. Wash again with 1 ml cold NIB+ and discard the supernatant. The final pellet volume should be ≤ 100 µl.

##### C. Chromatin Digestion

1. Resuspend the nuclei pellet in 300 µl of **1X** RE buffer.

   **CONTROL:** Keep aside 15 µl of this nuclei suspension as <u>nuclei control</u>. Store at -20 ºC until needed (end of day 3).
2. Centrifuge at 2,000 × g for 5 minutes at 4 ºC, discard supernatant.
3. Gently resuspend in 150 µl of 0.5% SDS (NOT ON ICE – the SDS would precipitate). Split into **<u>three</u> 2 ml safe-lock tubes** (~50 µl of nuclei suspension in each tube).
4. Incubate at 62 ºC for 5 minutes.
5. Quench SDS by adding 145 µl of water and 25 µl of 10% Triton X-100 to each tube.
6. Incubate at 37 ºC for 15 minutes.
7. Add 25 µl of RE buffer 10X and 60 U (6 µl) DpnII (The total volume in each tube should be ~251 µl at this point).
8. Mix by inversion.

9. Incubate **overnight** at 37 ºC, gently shaking (~60 rpm). To improve mixing, tubes should ideally be horizontal or almost horizontal. Seal lids with parafilm to make sure there is no spill.

#### DAY 2

10. Incubate at 62 ºC for 20 minutes, cool down at RT.

    **CONTROL:** Take 15 µl from each tube, and pool them in a single tube as digestion control. Store at -20 ºC until needed (end of day 3)

##### D. Affinity Labeling

1. To each tube, add:

    0.5 µl of 10 mM dTTP (15 µM final)

    0.5 µl of 10 mM dGTP (15 µM final)

    0.5 µl of 10 mM dCTP (15 µM final)

    5 µl of 1 mM biotin-14-dATP (15 µM final)

    43.5 µl of water

    4 µl of Klenow Fragment (40 U)

2. Mix gently by inversion (total volume in each tube should be ~300 µl at this point), incubate at **RT for 4 hours** (can be overnight). Seal lids with parafilm to make sure there is no spill, invert tube gently from time to time.

##### E. Proximity Ligation

3. To each tube add:

    660 µl of water

    120 µl of blunt end ligation buffer

    100 µl 10% Triton X-100

    20 µl T4 DNA ligase

4. Mix gently, incubate at 16 ºC 4 h, **overnight (O.N.)**, gently shaking (~60 rpm). To improve mixing, tubes should ideally be horizontal or almost horizontal. Seal lids with parafilm to make sure there is no spill.

#### DAY 3

5. Spin at 2,000 xg for 4 min at RT, discard supernatant.

##### F. DNA extraction

1. Resuspend in 750 µl of SDS lysis buffer.

   **CONTROLS**: do the same for the stored controls (250 ul SDS lysis buffer).
2. Add 25 µl proteinase K (20 mg/ml = 800 U/µl; 4 ul for controls).
3. Incubate 30 min at 55 ºC.
4. Add 32 µl 5 M NaCl (10 ul for controls), incubate at 65 ºC for 6 h or O.N. gently rotating/shaking. Seal lids with parafilm to make sure there is no spill.

#### DAY 4

5. Let tubes cool down, add 1 µl RNAseA 100 mg/ml (or 5 ul of a 20mg/ml solution).
6. Incubate 1 h at 37 ºC, shaking. gently
7. Add 10 µl proteinase K, incubate 2 more hours at 55 ºC, shaking.
8. Add 750 µl of phenol:chlorophorm:isoamyl alcohol 25:24:1.
9. Mix vigorously, spin at 13,000 xg for 5 minutes at RT.
10. Move aqueous phase to a new 2 ml tube.
11. Add 75 µl of 3M NaAc pH 5.2, mix.
12. Add 750 µl of cold isopropanol. Mix by inversion.
13. Incubate for 20 min at -20 ºC.
14. Spin for 15 minutes at full speed.
15. Wash pellet with 1 ml 75% ethanol, air dry pellet.
16. Resuspend in 45 µl 10 mM Tris-HCl.
17. <u>Combine DNA from the three tubes in a single tube</u>.
18. Measure concentration (Qubit BR).

19. Analyze samples profile in a 1% agarose gel (load 50-100 ng each of nuclei control, digestion control, final DNA). You expect the samples to look like in Figure 1. DNA in the nuclei control should be high molecular weight; after digestion, you should see a smear of smaller DNA fragments, and the size distribution should increase again after ligation in the final DNA.

20. Store DNA at -20 ºC until needed.

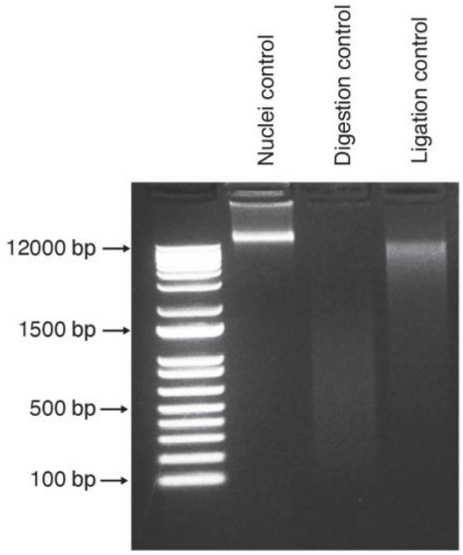


**Figure 1: DNA controls for Hi-C treatment**

#### Illumina library preparation

This section of the protocol describes a ligation-based Illumina library preparation, starting from the biotinylated Hi-C DNA obtained in the previous section. After step G, this protocol can be substituted with any other suitable protocol for the preparation of paired-end Illumina libraries, making sure to use streptavidin-coated beads to retain only relevant fragments (i.e. fragments representing Hi-C interactions). The sequence of adapters and primers used in this protocol is reported in Appendix III.

#### DAY 5

##### G. Removal of biotin from non-ligated ends and Shearing

1. Start with a total of 4 µg of DNA, divided in <u>three 1.33 µg aliquots</u> in **PCR tubes**.
2. To each aliquot, add:

   5 µl of 10X T4 polymerase buffer (NEB 2.1)

   0.5 µl of 10 mM dGTP

   1.75 µl of T4 DNA polymerase (5 U, 3000 U/ml)

   Water to 50 µl
3. Incubate 4 h at 20 ºC (in thermocycler).
4. Stop reaction by adding 1 µl of 0.5 M EDTA.
5. Add 79 µl of water to get to 130 µl.
6. Store at -20 ºC.

#### DAY 6

7. Shear to ~400 bp using a Covaris ultrasonicatior.

**NOTE:** On a Covaris ME220, using snap-cap (130 µl) tubes, the following conditions were used:

Program: Duration 30s

Peak power: 70

Duty Factor (%)

Cycles/Burst: 1000

Average power: 14

**H. Size selection**

1. Pool samples in a LoBind 1.5 ml tube.
2. Add 0.7 volumes of SPRI beads. Mix well by pipetting or vortexing.
3. Leave at room temperature for 10 minutes.
4. Move the tubes to the magnet, leave them there for 5 minutes or until the solution is clear.
5. Move supernatant to a clean tube.
6. Add 0.3 volumes of SPRI beads (for a total of 1 volume of beads), mix well, leave at room temperature for 15 minutes.
7. Move the tubes to the magnet, leave them there for 5 minutes or until the solution is clear.
8. Remove the supernatant.
9. Keeping the tubes on the magnet, add 500 µl of ethanol 75% to each tube.
10. Wait at least 30 seconds, then remove the ethanol. Repeat the wash.
11. Dry the tubes for 1-3 minutes at room temperature.
12. Resuspend well by pipetting in 50 µl of Tris 10 mM, pH 8.0.
13. Leave at room temperature for 5 minutes.
14. Put back on the magnet, wait for a couple of minutes or until the solution is clear. Move the supernatant to a clean tube.

**I. Biotin pulldown**

1. Vortex streptavidin C1 Dynabeads and transfer 15 µl to a 1.5 ml LoBind tube.
2. Add 300 µl of TWB buffer, mix by pipetting.
3. Incubate on a rocking platform 3 minutes at RT.
4. Put on magnet, remove supernatant.

5. Resuspend in 300 µl of TWB buffer, mix by pipetting, move to a new LoBind tube.
6. Incubate on a rocking platform 3 minutes at RT.
7. Put on magnet, remove supernatant.
8. Resuspend in 50 µl 2X BB, and add the 50 µl of size-selected DNA.
9. Incubate 30 min at RT, gentle rotation.
10. Put on magnet, discard supernatant (but keep in a tube just in case).
11. Resuspend in 500 µl of 1X BB. Move to a new LoBind tube.
12. Incubate 5 min at RT.
13. Put 5 min on magnet. Pipette up and down the supernatant once to make sure no beads are removed, then remove the supernatant.
14. Repeat wash in 500 µl 1X BB.
15. Resuspend in 100 µl TLE buffer, move to a new LoBind tube.
16. Put on magnet, remove supernatant.

#### DAY 7

**WGS Illumina library preparation**

##### J. End-repair

1. Add to the beads:

    42.5 µl of water

    5 µl of NEBNext End Repair 10X buffer

    2.5 µl of NEBNext End Repair enzyme mix
2. Incubate 30 minutes at **20 ºC** (RT).
3. Put on magnet, discard supernatant.
4. Add 500 µl TWB, incubate on a rocking platform 5 min at RT.
5. Put back on magnet, remove supernatant.
6. Repeat TWB wash.
7. Resuspend in 200 µl 1X BB, transfer to new LoBind tube.
8. Put on magnet, pipette up and down the supernatant once to make sure no beads are removed, remove supernatant.

9. Add 100 µl of **1X** NEB buffer 2, transfer to new LoBind tube.
10. Store at 4 ºC O.N. or proceed.
11. Put on magnet, remove supernatant.

**K. A-tailing**

1. Add:

    43 µl water

    5 µl NEB buffer 2

    1 µl 10 mM dATP

    1 µl Klenow Fragment exo-

2. Incubate for 30 min at **37 ºC** on heatblock or incubator.
3. Put on magnet, discard supernatant.
4. Add 500 µl TWB, incubate on a rocking platform 5 min at RT.
5. Put back on magnet, remove supernatant.
6. Repeat TWB wash.
7. Resuspend in 200 µl 1X BB, transfer to new LoBind tube.
8. Put on magnet, pipette up and down the supernatant once to make sure no beads are removed, remove supernatant.
9. Add 100 µl of TLE buffer, move to a new LoBind tube.

**L. Adapter ligation**

1. Put on magnet, remove supernatant
2. Add:

    23 µl water

    25 µl NEB Quick Ligase buffer

    0.5 µl of 10 µM PE1 adapter

    0.5 µl of 10 µM PE2 adapter

    1 µl NEB Quick ligase

3. Incubate 15 minutes at **20 ºC** (RT), then cool down to 10 ºC (ice).
4. Put on magnet, discard supernatant.

5. Add 500 µl TWB, incubate on a rocking platform 5 min at RT.
6. Put back on magnet, remove supernatant.
7. Repeat TWB wash.
8. Resuspend in 200 µl 1X BB, transfer to new LoBind tube.
9. Put on magnet, pipette up and down the supernatant once to make sure no beads are removed, then remove supernatant.
10. Add 200 µl of 10 mM Tris-HCl pH 8.0, move to a new LoBind tube.
11. Put back on magnet, remove supernatant.
12. Resuspend in 25 µl of 10 mM Tris-HCl pH 8.0.

#### DAY 8

##### M. Enrichment test

This is a test for the amplification efficiency of your library – the goal is to identify the number of cycles that would produce a sufficient amount of library in step N (depending on what are the requirements of your sequencing centre of choice, but we usually aim at ending up with at least 100-200 ng of library). At least 7-8 cycles of PCR are recommended, since the enrichment step also extends and complete the Illumina adapter (and assigns unique indices to the library).

1. Set up a PCR as follows:

   8.5 µl water

   1 µl 10 µM PE-PCR-F-i5 (indexed)

   1 µl 10 µM PE-PCR-R-i7 (indexed)

   12.5 µl KAPA HiFi HotStart 2X ReadyMix

   2 µl beads suspension

2. With the following program:

   98 ºC 2 minutes

   98 ºC 10 seconds

   62 ºC 30 second

   72 ºC 30 second. Repeat for 15 cycles

   72ºC 3 minutes

3. Take 4 µl out at 6, 9, 12.
4. Run the samples at 6, 9 and 12 cycles, and 4 µl of the final product (15 cycles) on a 2% gel.
5. Purify the rest of the PCR with SPRI beads.
    a. Measure volume of PCR product.
    b. Add 1.5 volumes of SPRI beads at room temperature.
    c. Incubate for 5-10 minutes.
    d. Transfer to the magnet and incubate for 5-10 minutes or until solution looks clear.
    e. Keep tubes on magnet and remove the supernatant.
    f. Add 750 ul of 75% Ethanol, wait 30 seconds and remove it.
    g. Repeat step f. and make sure to remove the ethanol completely.
    h. Air dry for 2 minutes.
    i. Resuspend beads in 10 µl of 10 mM Tris-HCl, pH 8.0.
    j. Incubate for 5-10 minutes.
    k. Transfer to the magnet and incubate for 5 minutes or until the solution is clear.
    l. Take the supernatant and transfer to a new tube.
6. Determine the optimal amount of cycles that would give you at least 100-200 ng of libraries over 6 PCR reactions (see step N), and that prevents over-amplification of the library (which would result in a shift up of the library distribution of the gel – see "15 cycles" in Figure 2). If there is a visible band for primer dimers at your chosen nr of PCR cycles, you might want to consider reducing the amount of primers in step N.

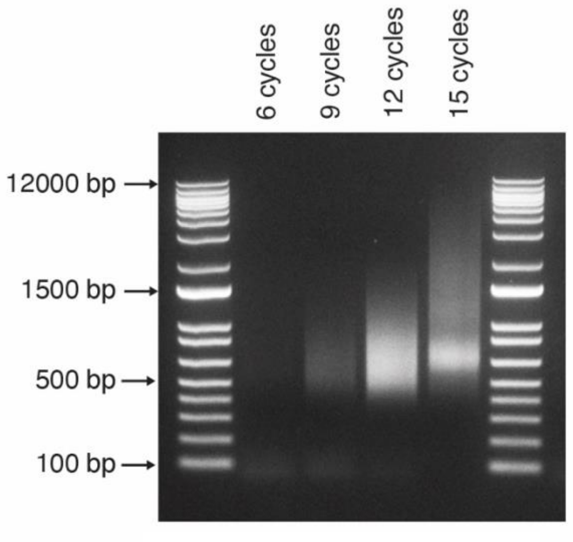


**Figure 2. Enrichment test**

##### N. Library Amplification

1. Do 6x PCR reactions with the same primers set used in the test, annealing 62 ºC, and the number of cycles chosen based on your gel and quantifications, sufficient to get 100-200 ng library in total (use 3 ul of library+ beads suspension from step L.12 as template for each PCR reaction).
2. Set up the PCR reactions as follows (x6):

   Water to 25 µl

   0.6-1 µl 10 µM PE-PCR-F-i5 (indexed)

   0.6-1 µl 10 µM PE-PCR-R-i7 (indexed)

   12.5 µl KAPA HiFi HotStart 2X ReadyMix

   3 µl beads suspension

   Note: the exact volume of primers will depend on the number of cycles you choose (see comments on step M.6)

#### DAY 9 (or continuation of day 8)

1. Pool PCRs, put on a magnet.
2. Keep supernatant in new tube. Add 12 µl of water to beads, keep at -20 ºC if needed (the original fragments are still bound to the beads).
3. Cleanup supernatant with 1.2V SPRI beads (180 µl).

   Note: verify volume of the supernatant.
4. Resuspend in 40 µl 10 mM Tris-HCl, pH 8.0.
5. Quantify on Qubit BR, run on Tapestation D1000 screentape.

#### Appendix I: Buffers and solutions

**PBS 1X (1L)**

- 8 g of NaCl (137 mM)
- 0.2 g of KCl (2.7 mM)
- 1.44 g of $Na_2HPO_4$ (10 mM)
- 0.24 g of $KH2PO_4$ (1.8 mM)

Dissolve the reagents listed above in 800 mL of H2O. Adjust the pH to 7.4 (or 7.2, if required) with HCl, and then add $H_2O$ to 1 L.

**Glycine 2M (20 ml)**

- 3 g in 20 ml water

**Percoll gradient (50 ml)**

- 4.2 g of sucrose (0.25 M)
- 50 µl of 1 M Tris-HCl pH 7.5 (1 mM)
- Add Percoll until 50 ml (95%)

**Nuclei Isolation Buffer (NIB) (100 ml), STORE AT 4 ºC**

- 1.5 ml of 1 M Tris HCl pH 8.0 (15 mM)
- 400 ul of 0.5 M EDTA pH 8.0 (2 mM)
- 8 ml of 1 M KCl (80 mM)
- 400 ul of 5 M NaCl (20 mM)

  **Add fresh**:
- 0.1% Triton X-100 (200 µl of 10% Triton X-100 for each 20 ml of NIB)
- 0.5 mM Spermine (200 µl of 50 mM spermine for each 20 ml of NIB)
- 0.1 mM PMSF (20 µl of 100 mM PMSF for each 20 ml of NIB)
- 1% beta-mercaptoethanol (20 µl of beta-mercaptoethanol for each 20 ml of NIB)
- 1% protease inhibitor cocktail (200 µl of protease inhibitor cocktail for each 20 ml of NIB)

**RE buffer 10X (5 ml)**

- 1 ml of 5 M NaCl (1 M)
- 2.5 ml of 1 M Tris-HCl, pH 7.9 (0.5 M)
- 500 µl of 1 M $MgCl_2$ (100 mM)
- 50 µl of 1 M DTT (10 mM)

**10X Blunt end ligation buffer (10 ml, aliquot)**

- 3 ml of 1 M Tris-HCl, pH 7.8 (300 mM)

- 1 ml of 1 M $MgCl_2$ (100 mM)
- 1 ml of 1 M DTT (100 mM)
- 1 ml of 10 mM ATP (1 mM)

**SDS lysis buffer (1 ml) PREPARE FRESH**

- 50 µl of 1 M Tris-HCl pH 8.0 (50 mM)
- 100 µl of 10% SDS (1%)
- 20 µl of 500 mM EDTA (10 mM)

**TLE buffer (10 ml)**

- 100 µl of 1 M Tris-HCl pH 8.0 (10 mM)
- 2 µl of 0.5 M EDTA (0.1 mM)

**2X BB (10 ml)**

- 100 µl of 1 M Tris-HCl pH 8.0 (10 mM)
- 20 µl of 0.5 M EDTA (1 mM)
- 4 ml of 5 M NaCl (2M)

**1X BB (25 ml)**

- 125 µl of 1 M Tris-HCl pH 8.0 (5 mM)
- 25 µl of 0.5 M EDTA (0.5 mM)
- 5 ml of 5 M NaCl (1 M)

**TWB buffer (50 ml)**

- 250 µl of 1 M Tris-HCl pH 8.0 (5 mM)
- 50 µl of 0.5 M EDTA (0.5 mM)
- 10 ml of 5 M NaCl (1 M)
- 25 µl of Tween-20 (0.05% v/v)

#### Appendix II: Reagents and equipment

For some of the reagents, the vendor we routinely used for this protocol is listed. However, equivalent products from alternative vendor will likely work as well. Basic lab equipment (pipettes, tips, conical tubes, etc.) are not listed. For handling nuclei, wide-bore pipette tips should be used – cutting the the tip of regular tips works as well.

| Reagents (Hi-C DNA preparation) | Vendor | Catalog number |
|---|---|---|
| Protease inhibitor | Millipore sigma | P9599-5ML |
| Phenylmethylsulfonyl fluoride (PMSF) | Millipore sigma | 10837091001 |
| Beta-mercaptoethanol | | |
| Triton X-100 | | |
| Spermine | | |
| Formaldehyde solution 37% | Millipore sigma | 47608-250ML-F |
| Glycine | | |
| Phosphate Buffered Saline (PBS) 10X | | |
| Miracloth | Millipore sigma | 475855-1R |
| Cheese cloth | | |
| Percoll | Millipore sigma | P1644-100ML |
| Sucrose | | |
| Tris HCl 1M (pH 7.5; 7.8; 7.9; 8.0) | | |
| EDTA 0.5 M pH8.0 | | |
| Dithiothreitol (DTT) | | |
| NaAc | | |
| KCl | | |
| NaCl | | |
| $MgCl_2$ | | |
| DpnII (10,000 units/ml) | NEB | R0543L |
| Klenow fragment (10 U/μL) | ThermoFisher Sci. | EP0052 |
| dNTPs (100 mM, individual) | ThermoFisher Sci. | 10297018 |
| Biotin-14-dATP | Jena Bioscience | NU-835-BIO14-S |
| T4 DNA ligase (2,000,000 units/ml) | NEB | M0202M |
| Phenol:chloroform:isoamylalcohol | ThermoFisher Sci. | 15593031 |
| RNAseA | | |
| Proteinase K (800 units/ml) | NEB | P8107S |
| Isopropanol | | |
| Ethanol 75% | | |
| Qubit assay tubes | ThermoFisher Sci. | Q33252 |
| Qubit assay BR kit | ThermoFisher Sci. | Q32853 |

| Reagents (Illumina library prep) | Vendor | Catalog number |
|---|---|---|
| T4 DNA polymerase (3,000 units/ml) | NEB | M0203L |
| SPRI beads (Rohland and Reich, 2012) | | Or AMPure XP |
| Dynaeabd MyOne Streptavidin C1 | ThermoFisher Sci. | |
| Klenow fragment exo- (5,000 units/ml) | NEB | M0212L |
| Tween-20 | | |
| NEB Quick ligation kit | NEB | M2200L |
| NEB End Repair module | NEB | E6050L |
| KAPA HiFi HotStart ReadyMix | Roche | 7958935001 |
| Qubit assay HS kit | ThermoFisher Sci. | Q32854 |
| Tapestation loading tips | Agilent Technologies | 5067-5599 |
| Tapestation D1000 reagent | Agilent Technologies | 5067-5583 |
| Tapestation D1000 screentape | Agilent Technologies | 5067-5582 |

**Consumable (beyond standard lab consumables)**

| | | |
|---|---|---|
| 1.5 ml DNA LoBind tubes | Eppendrof | 0030108051 |
| Covaris AFA tubes | | |

**Equipment (beyond standard lab equipment)**

| | | |
|---|---|---|
| Refrigerated centrifuge, 50 ml tubes | | |
| Refrigerated centrifuge, 1.5 ml tubes | | |
| Rotary incubator | | |
| Shaking incubator (16 ºC, 37 ºC, 65 ºC) | | |
| Qubit fluorometer | ThermoFisher Scien. | |
| Covaris ultrasonicator | Covaris | |
| Tapestation | Agilent Technologies | |
| Magnet for 1.5 ml tubes | ThermoFisher Scien. | 12321D |
| Magnet for PCR tube strips/plates | ThermoFisher Scien. | |

12321D
Magnet for PCR tube strips/plates

#### Appendix III: Adapters and primers

##### ADAPTERS

PE1 and PE2 adapters are ordered as individual oligos and then annealed

| | |
|---|---|
| PE1 adapter_top | 5′ TTTCCCTACACGACGCTCTTCCGATC*T 3′ |
| PE1 adapter_bottom | 5′ P-GATCGGAAGAGCGTCGTGTAGGGAAA 3′ |
| PE2 adapter_top | 5′ GTTCAGACGTGTGCTCTTCCGATC*T 3′ |
| PE2 adapter_bottom | 5′ P-GATCGGAAGAGCACACGTCTGAAC 3′ |

* = phosphorothioate bond
P = 5' phosphorylation

**Adapter annealing**
Resuspend oligos to 100 μM in 10 mM TRIS pH8.0.

| | |
|---|---|
| Top stand oligo (100 μM) | 5 μl |
| Bottom strand oligo (100 μM) | 5 μl |
| 10x annealing buffer | 5 μl |
| water | 35 μl |

in a thermal cycler, bring at 95 °C for 5 minutes, and then ramp down to room temperature at 0.1 °C/second. The resulting concentration of the adapter is ~10μM.

**10X annealing buffer**
100 mM Tris pH8.0
0.5 M NaCl

**PCR primers**
These primers complete the adapters and add i7 and i5 indices (usually 6-8 unique bp sequences, here shown as XXXXXXXX)

PE-PCR-F-I5
5′ AATGATACGGCGACCACCGAGATCTACACXXXXXXXXACACTCTTTCCCTACACGACGCT 3′

PE-PCR-R-I7-INDEX193
5′ CAAGCAGAAGACGGCATACGAGATXXXXXXXXGTGACTGGAGTTCAGACGTGTGCTCTT 3′
